# Node persistence from topological data analysis reveals changes in brain functional connectivity

**DOI:** 10.1101/2025.05.23.655816

**Authors:** Madhumita Mondal, Yasharth Yadav, Jürgen Jost, Areejit Samal

## Abstract

Large-scale analyses of brain functional connectivity can uncover disruptions in regional activity and connectivity that are commonly associated with neurological disorders or cognitive decline associated with healthy aging. In our study, we employ persistent homology (PH), a prominent tool in topological data analysis (TDA), to investigate changes in resting-state functional connectivity in healthy aging and autism spectrum disorder (ASD). We analyze functional connectivity changes across three distinct scales: (a) global scale (brain-wide changes), (b) mesoscopic scale (resting-state network-level changes), and (c) local scale (region-level changes). At the local scale, we introduce node persistence, a novel and scalable PH-based measure that detects brain regions with significant differences in healthy aging or ASD. Notably, these regions overlap with regions whose non-invasive stimulation improves motor function in the elderly or alleviates ASD symptoms, suggesting the utility of node persistence in identifying clinically relevant brain regions affected by aging and ASD.

## INTRODUCTION

Brain functional connectivity analysis is a crucial aspect of neuroscience that investigates how different brain regions interact and communicate. Specifically, it examines temporal correlations between neuronal activity across spatially distinct brain regions, revealing activation patterns that occur during cognitive tasks or at rest [1]. Such analyses typically employ neuroimaging techniques such as functional magnetic resonance imaging (fMRI), electroencephalography (EEG), or magnetoencephalography (MEG). Among these, fMRI is particularly prominent due to its ability to capture spontaneous neuronal activity by measuring fluctuations in blood-oxygen-level-dependent (BOLD) signals. Pairwise correlations between BOLD signal time series across different brain regions [2–4] can reveal functional connectivity networks underlying the brain. Interestingly, spatially distant brain regions demonstrate synchronized activity even in the absence of external cognitive tasks. This phenomenon is known as resting-state functional connectivity, and is estimated from resting-state fMRI (rs-fMRI) data [5–7]. Coherent patterns of this activity across anatomically distinct regions are organized into resting-state networks (RSNs), which reflect the intrinsic functional architecture of the brain. These intrinsic functional networks provide a foundation for understanding large-scale brain organization in both health and disease [5, 7–10]. In this study, we focus on studying functional connectivity alterations in two distinct biological processes, age-related cognitive decline, as well as atypical neurodevelopment associated with autism spectrum disorder (ASD).

As the global population ages, uncovering the neural correlates of age-related cognitive decline has become increasingly important. Neuroimaging techniques such as fMRI have been instrumental in advancing our understanding of healthy brain aging [11, 12]. Many studies have identified functional changes in regions such as the prefrontal, medial temporal, and parietal cortices, which are essential for maintaining cognitive performance in elderly individuals [11, 13]. In parallel, neuroimaging tools have been applied to investigate neurodevelopmental disorders, most notably ASD. A substantial amount of research has investigated the pathophysiology and neurobiology of ASD, providing critical insights into the structural and functional development in ASD relative to typical development [14]. ASD refers to a broad range of neurodevelopmental conditions [15], typically characterized by difficulties in social interaction, verbal or non-verbal communication, restrictive or repetitive behaviors, [14, 15] often accompanied by varying levels of cognitive, motor, or memory impairments [16–18]. The prevalence of ASD is increasing worldwide [19, 20], and while early diagnosis is crucial for effective intervention, it remains equally important to ensure diagnostic reliability and accuracy [21]. Together, these findings underscore the role of neuroimaging techniques, particularly fMRI, in elucidating the neural mechanisms underlying both healthy aging and ASD [22–24]. However, the high complexity and dimensionality of fMRI data have driven the need for advanced analytical tools capable of capturing altered functional connectivity beyond conventional methods. Topological data analysis (TDA) offers a robust, multiscale framework for extracting meaningful features from such high-dimensional datasets [25].

By incorporating principles from algebraic topology and computational geometry, TDA characterizes the inherent shape of high-dimensional datasets [26–28]. In this study, we focus on persistent homology (PH), a central tool in TDA that captures topological features or “holes” in the dataset at multiple scales. These features include connected components (*H*_0_), loops (*H*_1_), and voids (*H*_2_), which correspond to zero, one, and two dimensional holes, respectively. These topological features can then be compactly summarized using representations such as persistence barcodes, persistence diagrams, or persistence landscapes, each capturing the evolution of features across multiple scales [29– 31]. Unlike traditional graph-theoretic or network-based methods, which often rely on ad hoc threshold parameters to define connectivity, PH operates across multiple scales, eliminating the need for arbitrary parameter selection. This multiscale framework enables a more robust and unbiased characterization of topological features in complex datasets [31, 32]. PH has found widespread application across diverse scientific domains, including biology [33–36], finance [37–40], physics [41–43], machine learning [44–46], as well as image and signal processing [47, 48]. In particular, PH has found prominent applications in neuroscience due to its ability to encapsulate the multiscale structure of brain connectivity [49–61]. It has been shown to effectively distinguish typical and clinically impaired brain states, providing valuable insights into neurological conditions like ASD [55, 56], Parkinson’s disease [58], Alzheimer’s disease [54, 57], and age-related cognitive impairment [59].

While PH is well-suited for summarizing the global topological structure of complex datasets, it lacks the ability to provide local topological information [30, 31]. This limitation arises because homology groups are computed on simplicial complexes constructed from the entire dataset, thereby obscuring node-level details. Furthermore, the representatives of homology classes, that is cycles or holes, are inherently non-unique [31, 62]. Nevertheless, understanding how specific brain regions influence global topological changes is crucial in functional connectivity analysis, especially in clinical applications where local alterations often contribute to global functional disruptions [5, 8, 10]. To address this limitation, several studies have investigated the potential for extracting local topological information from PH [46, 49, 50, 63–65]. In particular, Lord *et al*. [50] showed that in healthy individuals, topologically central nodes in the persistence scaffold, which summarize one-dimensional holes, may support functional integration across brain modules. Additionally, Liang *et al*. [55] constructed simplicial complexes at specific thresholds and found significantly fewer one-dimensional holes within certain brain regions in the ASD group compared to the typically developing (TD) controls. However, as data size and dimensionality increase, these methods tend to become increasingly computationally expensive. As a result, local techniques based on PH have seen limited applications in large-scale neuroimaging datasets, highlighting the need for more scalable and efficient methodological advancements.

The primary goal of our work is to develop efficiently computable PH-based metrics and to utilize them to identify alterations in the brain functional connectivity in healthy aging and ASD, and to link them with cognition and behaviour. To this end, we utilized functional connectivity (FC) matrices of 225 subjects from the MPI-LEMON dataset [66, 67] and 820 subjects from the ABIDE-I dataset [68, 69]. First, we computed PH-based measures at three distinct scales, namely global, mesoscopic, and local, to compare healthy young and healthy elderly individuals in the MPI-LEMON dataset, and ASD and TD individuals in the ABIDE-I dataset. At the local or region-level scale, we introduced two novel PH-based metrics, *node persistence* and *node frequency*, which quantify the contribution of a node to one-dimensional homological features. These measures are computationally more efficient than existing local PH-based measures, making them suitable for large-scale brain connectivity analysis. Fig. 1 presents an overview of PH-based measures. Second, we utilized Neurosynth meta-analysis [70, 71] to identify the cognitive domains associated with those brain regions that exhibit significant group differences in our proposed local topological measures. Third, we conducted a correlation analysis to examine the relationship between PH-based measures and phenotypic test scores of individuals in the MPI-LEMON dataset, as well as clinical symptom severity scores for individuals with ASD. Fourth, we compared the regions with significant between-group differences in local topological measures against clinically relevant regions documented in existing non-invasive brain stimulation (NIBS) studies. Fifth, we evaluated the effectiveness of our proposed metrics by comparing them with an existing PH-based method for detecting region-level topological changes.

**FIG. 1.**
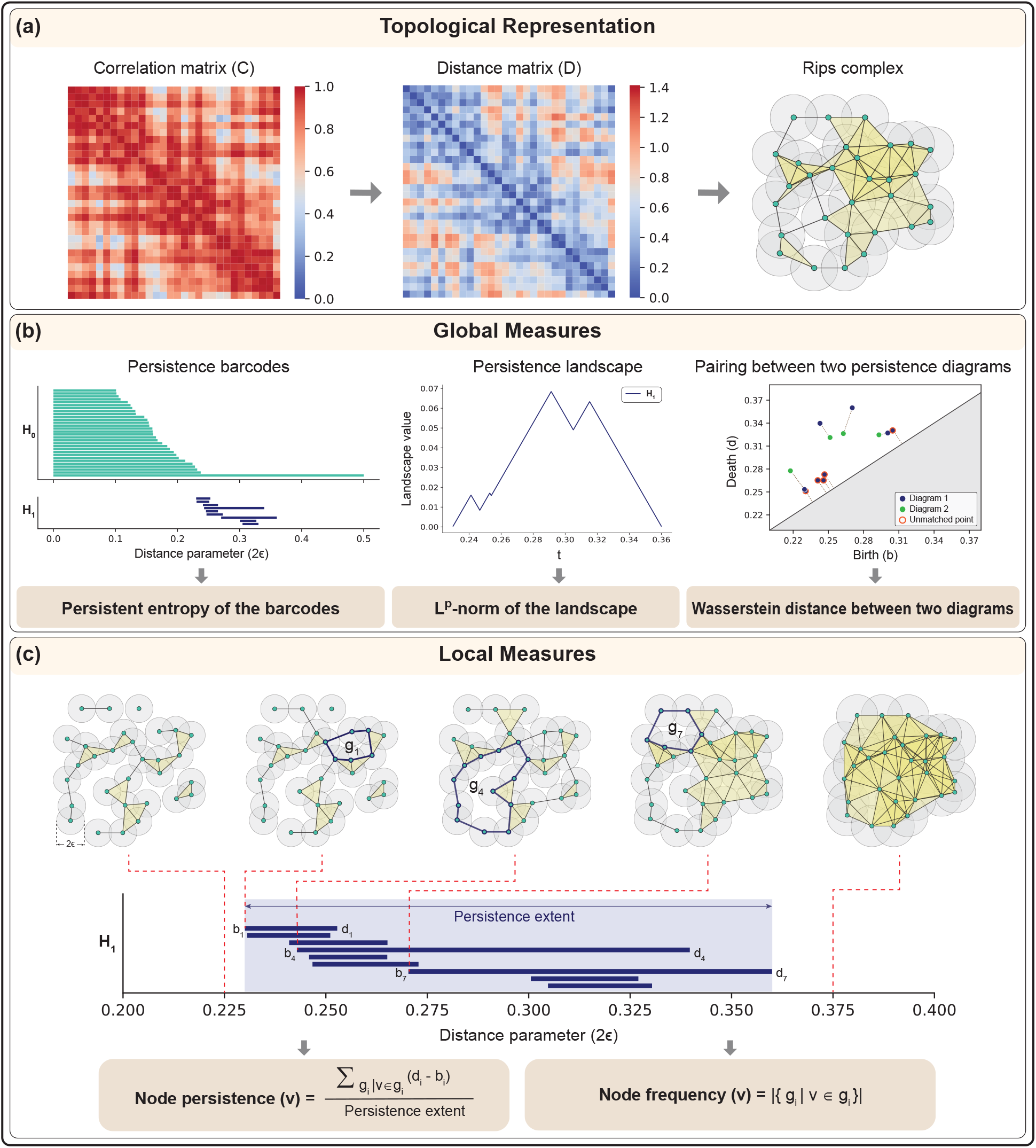
Schematic overview of persistent homology based characterization of functional connectivity. (a) Topological representation: from the functional connectivity matrix, which is a type of correlation matrix, the distance matrix 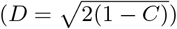 was constructed. Next, this distance matrix was used to build a Rips complex. Note that the Rips complex shown may not reflect the exact spatial embedding of the nodes from the distance matrix. (b) Global measures: we computed persistent entropy, which quantifies the Shannon entropy of the persistence barcodes, and the *L*^1^-norm and *L*^2^-norm, which capture the overall magnitude of the persistence landscape, to investigate brain-wide and resting-state network-level changes in functional connectivity. Next, we compared the 1-Wasserstein, 2-Wasserstein, and bottleneck distances between intra-group and inter-group persistence diagrams. To compute these distances, points in the diagrams are matched pairwise, or with the diagonal if no suitable pairing is possible, and the distances are then calculated based on the lengths of the connecting lines. (c) Local measures: we computed node persistence and node frequency to characterize region-level changes in functional connectivity. These measures quantify the influence of a node (*v*) on one-dimensional homological features (*H*_1_). Rips complexes corresponding to five different distance parameters (2*ϵ*) are shown, where *ϵ* denotes the radius of the balls centered at each point; *g*_*i*_ denotes a representative cycle in *H*_1_ with birth *b*_*i*_ and death *d*_*i*_.

## RESULTS

In this study, we applied topological data analysis (TDA) to investigate changes in resting-state functional connectivity associated with healthy aging and autism spectrum disorder (ASD). These changes were characterized using topological measures based on PH across three spatial scales: (a) global or brain-wide changes, (b) mesoscopic or resting-state network (RSN)-level changes, and (c) local or region of interest (ROI)-level changes. We acquired functional connectivity (FC) matrices from the MPI-LEMON dataset for the healthy aging investigation and the ABIDE-I dataset for the ASD investigation. The FC matrices in the MPI-LEMON dataset correspond to 153 healthy young and 72 healthy elderly individuals, while those in the ABIDE-I dataset correspond to 395 individuals with ASD and 425 TD individuals. These FC matrices were derived in earlier studies conducted by some of us [67, 69] from publicly available resting-state fMRI (rs-fMRI) scans. Each FC matrix is a 200*×*200 square matrix representing pairwise Pearson correlations between the 200 ROIs, as specified by the Schaefer atlas [72]. Next, we filtered the FC matrices to retain only positive correlations and converted them to ultrametric distance matrices [73]. Our analysis focused on positive correlations, as they have a primary and central role in the structure and organization of brain functional connectivity networks [74]. Finally, we constructed a filtration of Vietoris-Rips (Rips) complexes on each distance matrix and computed PH-based measures to extract multiscale topological information from the FC matrices (see Fig. 1(a)). A detailed description of the FC matrices, Rips complex construction, and topological measures is provided in the Methods section.

### Brain-wide differences in functional connectivity

We determined brain-wide changes in brain functional connectivity between the groups of individuals using three global topological measures derived from PH, namely persistent entropy of the persistence barcodes associated with *H*_0_, *H*_1_, and *H*_2_ [75], *L*^1^-norm and *L*^2^-norm of the persistence landscape associated with *H*_1_ [76] (see Fig. 1(b)). To evaluate statistical differences between the groups, a two-tailed two-sample t-test was conducted [77].

Fig. 2(a) shows violin plots comparing persistent entropy, *L*^1^-norm, and *L*^2^-norm between the young and elderly groups in the MPI-LEMON dataset. In the young group, mean persistent entropy is significantly higher (*p <* 0.001), suggesting that the persistence of its features is more evenly distributed than in the elderly group. Furthermore, we found that the young group shows a higher mean *L*^1^-norm (*p <* 0.01) and *L*^2^-norm (*p <* 0.001) of the persistent landscape compared to the elderly group. This indicates that one-dimensional holes remain more persistent in the young group. Fig. 2(b) shows violin plots comparing persistent entropy, *L*^1^-norm, and *L*^2^-norm between the ASD and TD groups in the ABIDE-I dataset. We found that the mean persistent entropy is significantly higher in the ASD group compared to the TD group (*p <* 0.001). In contrast, the mean *L*^1^-norm and *L*^2^-norm are significantly lower in the ASD group compared to the TD group (*p <* 0.001 for both).

**FIG. 2.**
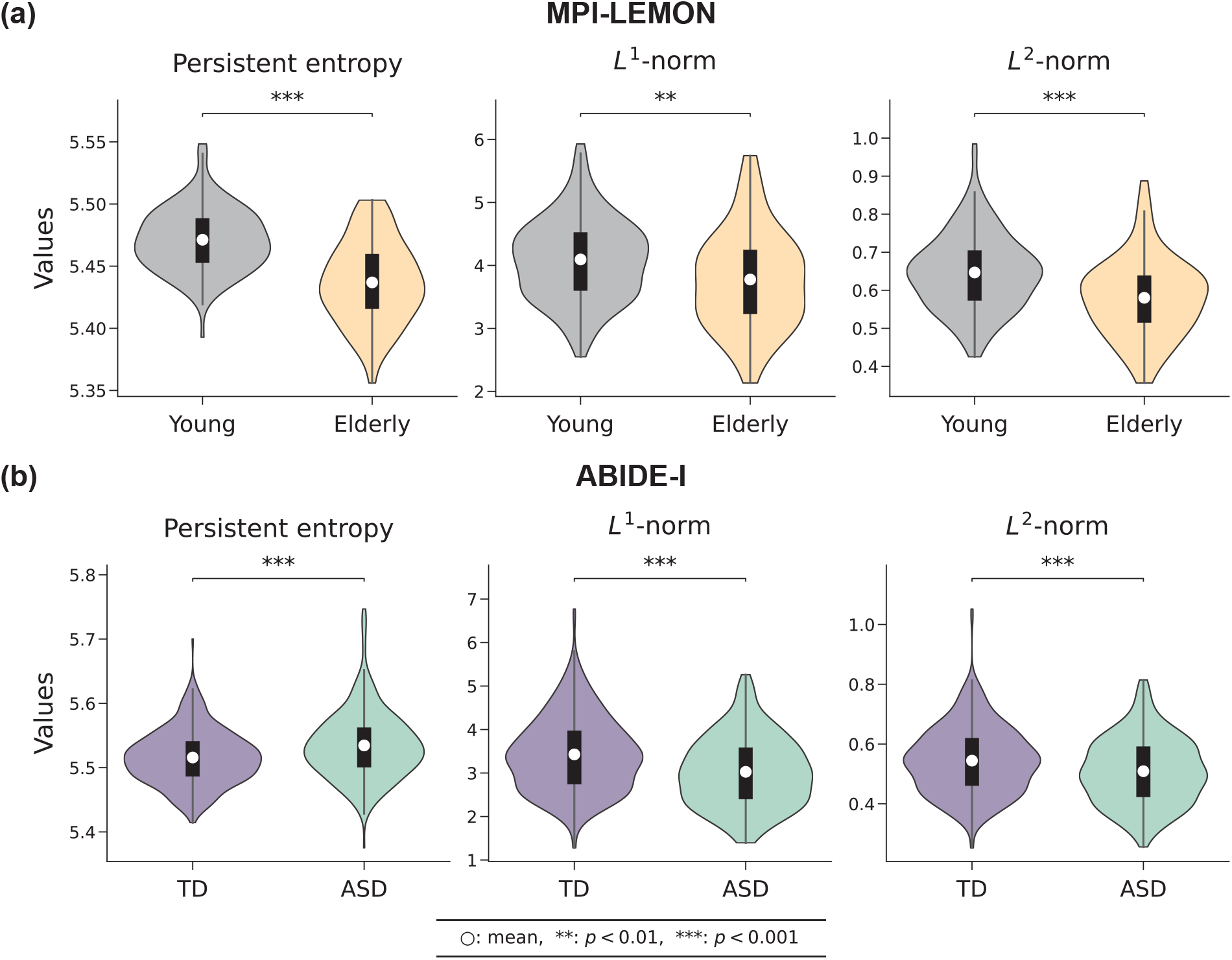
Brain-wide differences between the groups as identified by three global measures: persistent entropy of the persistence barcodes, *L*^1^-norm and *L*^2^-norm of the persistent landscape. (a) MPI-LEMON dataset: violin plots corresponding to 153 young and 72 elderly individuals across three global measures. The mean values of all three measures are significantly higher (*p <* 0.01) in the young group compared to the elderly group. (b) ABIDE-I dataset: violin plots corresponding to 425 typically developing (TD) individuals and 395 individuals with autism spectrum disorder (ASD) across three global measures. Average persistent entropy is significantly higher (*p <* 0.001) in the ASD group than the TD group; however, average *L*^1^-norm and *L*^2^-norm are significantly lower (*p <* 0.001) in the ASD group.

The group-wise comparisons presented in Fig. 2 correspond to Rips complexes constructed from FC matrices with positive correlations only. We also constructed Rips complexes from FC matrices containing the full range of Pearson correlations and found that the group differences in the three global measures remained consistent (*p <* 0.05) with those obtained using only positive correlations in both the MPI-LEMON and ABIDE-I datasets. Supplementary Fig. S1 displays violin plots comparing the three global measures across groups in the MPI-LEMON and ABIDE-I datasets, considering the full range of Pearson correlations from the FC matrices. The findings imply that positive correlations significantly impact overall brain functional connectivity, as the global outcomes using all correlations closely resemble those obtained solely from positive correlations. The result also highlights that positive correlations are sufficient to reflect the overall functional organization of the brain.

In addition to comparing global topological measures between groups, we compared the 1-Wasserstein distance, 2-Wasserstein distance, and bottleneck distance between intra-group and inter-group persistence diagrams for both datasets [32]. In this scenario, we conducted a one-tailed two-sample t-test to evaluate statistical differences between them, as our null hypothesis is that intra-group distances would be less than inter-group distances. Supplementary Figures S2 and S3 illustrate the corresponding violin plots based on positive correlations and all correlations, respectively. For all three distance measures, inter-group distances are found to be significantly higher than intra-group distances across both datasets. Supplementary Table S1 reports the average values for each group along with the corresponding p-values between them for the measures: persistent entropy, *L*^1^-norm, and *L*^2^-norm. For the 1-Wasserstein distance, 2-Wasserstein distance, and bottleneck distance, it presents the intra-group and inter-group average values as well as the p-values between them. Supplementary Table S1 also includes the results based on both positive correlations and all correlations from the FC matrices.

### RSN-level differences in functional connectivity

RSNs are critical for understanding the intrinsic functional organization of the brain without the influence of external tasks [10]. In this study, we focused on seven well-known RSNs defined according to the Schaefer atlas, specifically: *visual network* (29 ROIs), *somatomotor network* (35 ROIs), *dorsal attention network* (26 ROIs), *salience/ventral attention network* (22 ROIs), *limbic network* (12 ROIs), *control network* (30 ROIs), and *default network* (46 ROIs) [10, 72]. We examined each of the seven RSNs separately by analyzing the submatrices of the FC matrix that include entries corresponding to nodes or ROIs within that RSN. These submatrices were then transformed into distance matrices (see Methods section). As a result, seven unique distance matrices were created for each participant, each aligned with one of the seven RSNs. The size of each distance matrix depends on the number of nodes contained in the respective RSN. We assessed changes in brain functional connectivity at the RSN-level between the groups of individuals by applying persistent entropy of the persistence barcodes associated with *H*_0_, *H*_1_, and *H*_2_, *L*^1^-norm, and *L*^2^-norm of the persistence landscape associated with *H*_1_. A two-tailed two-sample t-test was conducted to evaluate statistical differences between the groups, and False Discovery Rate (FDR) correction was applied separately for each measure.

For the MPI-LEMON dataset, violin plots comparing persistent entropy, *L*^1^-norm, and *L*^2^-norm across the young and elderly groups for each RSN are presented in Fig. 3(a). In the visual network, only the *L*^1^-norm and *L*^2^-norm show significant differences (*p <* 0.05, FDR-corrected) between the young and elderly groups. In the somatomotor, dorsal attention, salience/ventral attention, and default networks, all three measures show significant differences (*p <* 0.05, FDR-corrected) between the groups. In the limbic network, none of the three measures shows significant differences. In the control network, only persistent entropy shows a significant difference between the groups. For the ABIDE-I dataset, Fig. 3(b) presents violin plots comparing persistent entropy, *L*^1^-norm, and *L*^2^-norm across the ASD and TD groups for each RSN. In the somatomotor, salience/ventral attention, and default networks, all three measures show significant differences (*p <* 0.05, FDR-corrected) between the groups. In the visual, dorsal attention, limbic, and control networks, none of the three measures show significant differences.

**FIG. 3.**
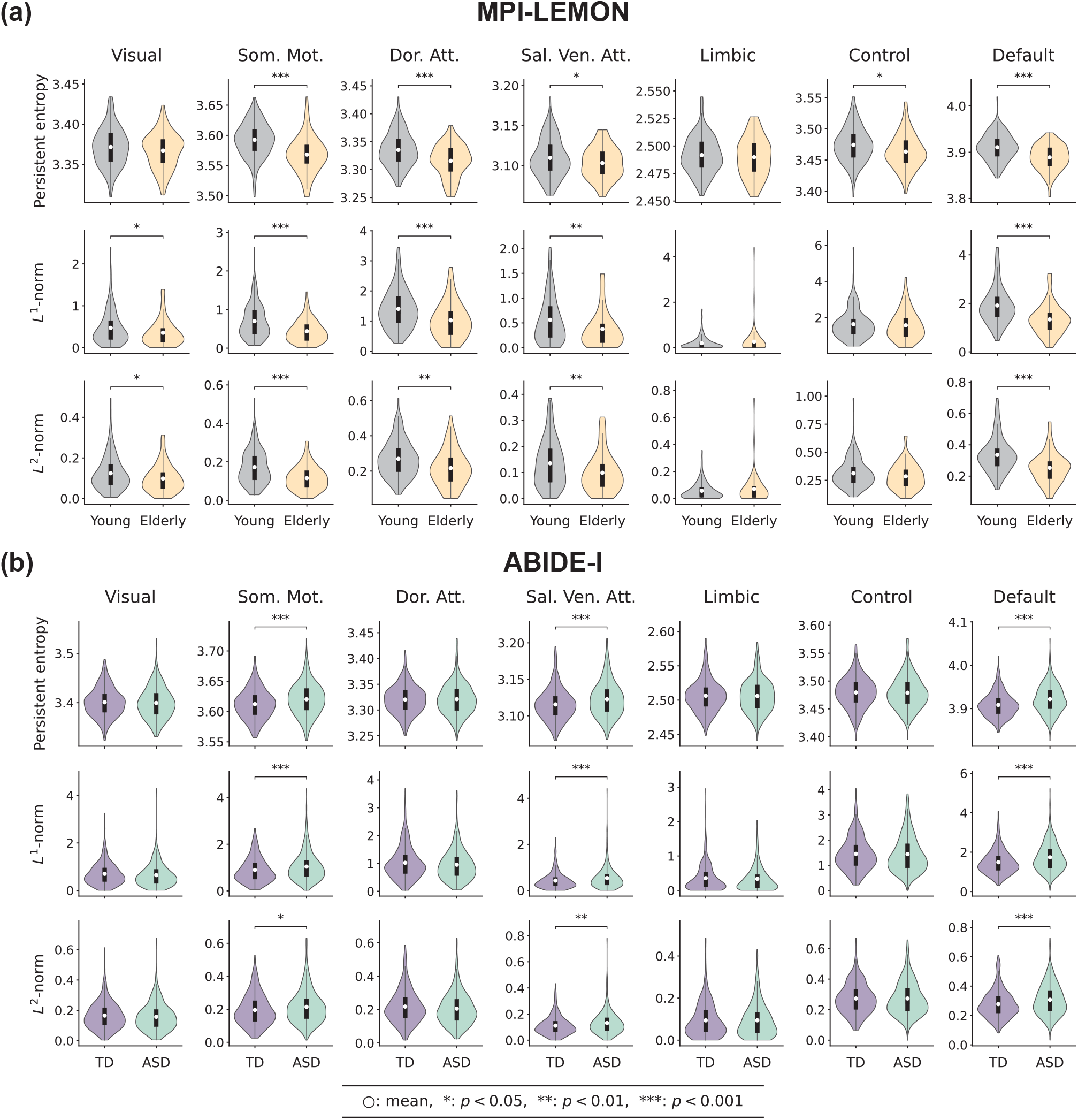
Resting-state network(RSN)-level differences between the groups as identified by three global measures: persistent entropy of the persistence barcodes, *L*^1^-norm and *L*^2^-norm of the persistent landscape. Each row corresponds to a given measure, and each column corresponds to a given RSN. (a) MPI-LEMON dataset: violin plots corresponding to 153 young and 72 elderly individuals across seven RSNs for each global measure are shown. For the somatomotor (Som. Mot.), dorsal attention (Dor. Att.), salience/ventral attention (Sal. Ven. Att.) and default networks, the mean values of all the three measures are significantly higher (*p <* 0.05, FDR-corrected) in the young group compared to the elderly group. For the visual network, only the mean values of the *L*^1^-norm and *L*^2^-norm are significantly higher (*p <* 0.05, FDR-corrected) in the young group. For the control network, only the mean of persistent entropy in the young group exhibits a significantly higher (*p <* 0.05, FDR-corrected) value than the elderly group. No significant differences are observed between the groups in the limbic network for any of the three measures. (b) ABIDE-I dataset: violin plots corresponding to 425 typically developing (TD) individuals and 395 individuals with autism spectrum disorder (ASD) across seven RSNs for each global measure are shown. For the somatomotor (Som. Mot.), salience/ventral attention (Sal. Ven. Att.), and default networks, the mean values of all three measures are significantly lower (*p <* 0.05, FDR-corrected) in the ASD group compared to the TD group. For visual, dorsal attention (Dor. Att.), limbic, and control networks, none of the three measures exhibit significant differences between the TD and ASD groups.

Additionally, we compared the 1-Wasserstein distance, 2-Wasserstein distance, and bottleneck distance between the intra-group and inter-group persistence diagrams for both datasets. To assess the significance of these differences, we performed one-tailed two-sample t-tests. Supplementary Fig. S4 presents violin plots illustrating the differences between intra-group and inter-group distances across both datasets. Supplementary Table S2 provides the average values for each group along with the FDR-corrected p-values between the groups for measures: persistent entropy, *L*^1^-norm, and *L*^2^-norm, corresponding to each RSN. For 1-Wasserstein distance, 2-Wasserstein distance, and bottleneck distance, it presents the intra-group and inter-group average values along with the FDR-corrected p-values corresponding to each RSN.

### ROI-level differences in the functional connectivity

In the first subsection, using whole-brain FC matrices, we showed that global topological measures based on PH can effectively differentiate between young and elderly groups in the MPI-LEMON dataset, as well as ASD and TD groups in the ABIDE-I dataset. In the second subsection, we extended this analysis to the RSN-level by applying the same topological measures to RSN-specific FC matrices. This enabled us to identify the RSNs where group-level differences in functional connectivity are concentrated. In this subsection, we focus on the 200 regions, as defined by the Schaefer atlas, to investigate ROI-level differences of the brain functional connectivity. To achieve this, we proposed two novel topology-based local measures: node persistence and node frequency (see Fig. 1(c) and Methods section). Node persistence quantifies the topological importance of a node by evaluating the duration of its involvement in one-dimensional holes during the filtration process. A higher node persistence value indicates that the node is consistently involved in more persistent one-dimensional holes, suggesting a key role in maintaining the loop structures of the simplicial complex. On the other hand, node frequency measures how many distinct one-dimensional holes a node belongs to, irrespective of their persistence duration throughout the filtration. We computed node persistence and node frequency across the 200 ROIs corresponding to each subject. A two-tailed two-sample t-test was used to evaluate group-level differences, and FDR correction was applied to adjust for multiple comparisons.

Based on node persistence, we identified several ROIs with significant between-group differences in both datasets. In the MPI-LEMON dataset, 108 ROIs exhibit significant differences (*p <* 0.05, FDR-corrected) between the young and elderly groups. Among the seven RSNs, the ROIs are categorized as: visual (12), somatomotor (22), dorsal attention (17), salience/ventral attention (11), limbic (2), control (15), and default (29) networks. All of these ROIs, except for one in the limbic network (RH Limbic TempPole 1), show higher node persistence values in young individuals compared to elderly individuals. In the ABIDE-I dataset, 27 ROIs exhibit significant differences (*p <* 0.05, FDR-corrected) between the ASD and TD groups. The ROIs are distributed as: visual (2), somatomotor (9), dorsal attention (1), salience/ventral attention (1), limbic (3), control (4), and default (7) networks. All the ROIs show higher node persistence values for individuals with ASD. Figures 4(a) and (b) depict the ROIs with statistically significant between-group differences (*p <* 0.05, FDR-corrected) based on node persistence for the MPI-LEMON and ABIDE-I datasets, respectively.

**FIG. 4.**
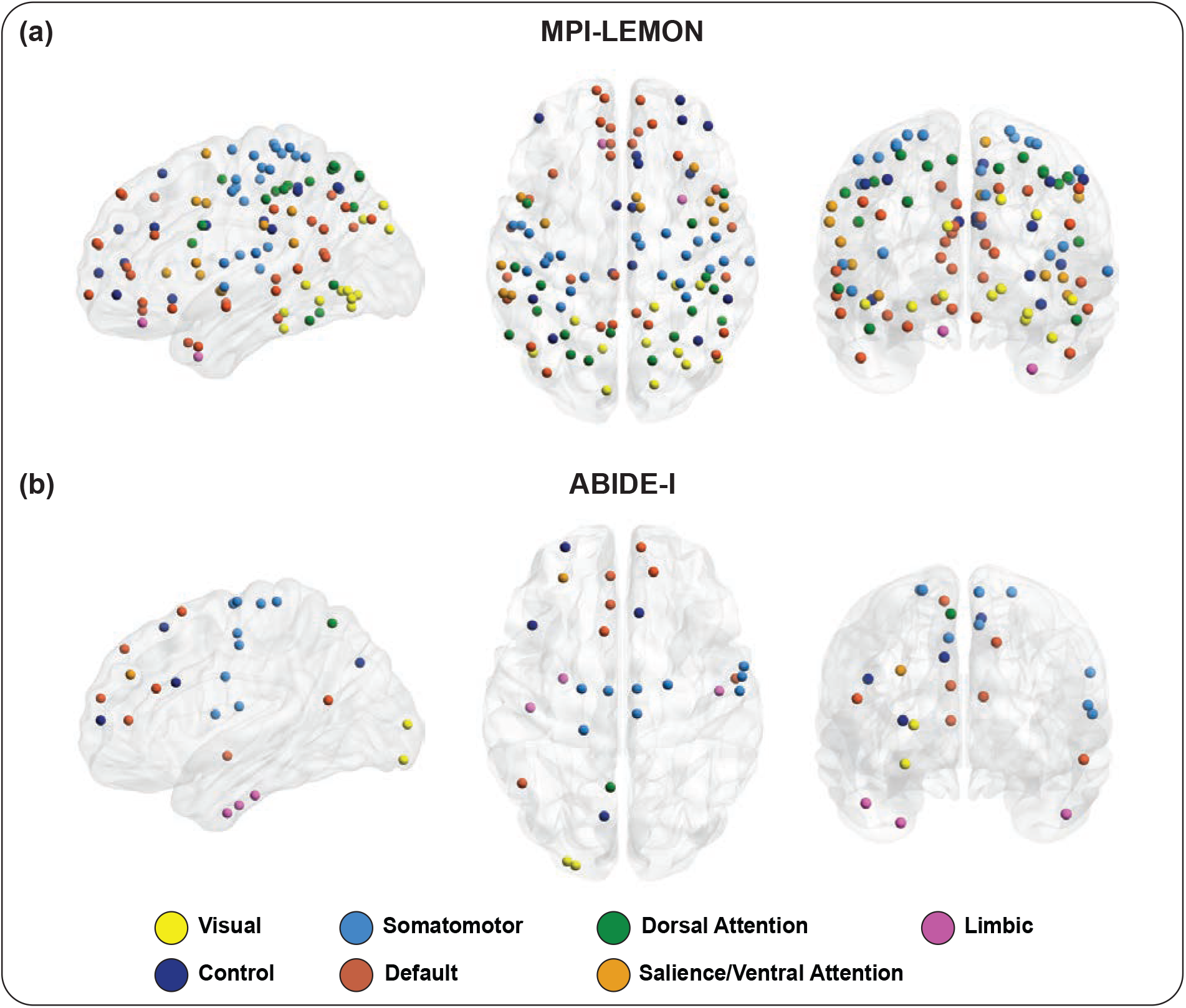
Visual representation of brain regions with significant between-group differences in node persistence (*p <* 0.05, FDR-corrected). (a) MPI-LEMON dataset: 108 regions with significant differences in node persistence between the healthy young and healthy elderly groups. For every region, young participants show higher node persistence compared to the elderly participants, except one region in limbic network (RH Limbic TempPole 1). (b) ABIDE-I dataset: 27 regions with significant differences in node persistence between the autism spectrum disorder (ASD) and typically developing (TD) groups. All regions reveal higher node persistence for ASD participants relative to TD participants. Each brain region is assigned to one of the seven resting-state networks (RSNs) as defined by the Schaefer atlas. The regions are colored according to their respective RSNs, as detailed in the figure legend. The visualization was generated using BrainNet Viewer [78]. Supplementary Table S3 lists the significantly different ROIs identified via node persistence across both datasets.

In contrast, node frequency reveals 39 and 35 ROIs with significant between-group differences (*p <* 0.05, FDR-corrected) for the MPI-LEMON and ABIDE-I datasets, respectively (see Supplementary Fig. S5). In the MPI-LEMON dataset, 39 ROIs are distributed as follows: visual (3), somatomotor (6), dorsal attention (9), salience/ventral attention (4), limbic (2), control (3), and default (12) networks. Moreover, the ROIs identified through node frequency are a subset of those identified by node persistence. In the ABIDE-I dataset, 35 ROIs are distributed in six RSNs as follows: visual (5), somatomotor (11), dorsal attention (2), salience/ventral attention (5), control (5), and default (7) networks. Moreover, 21 out of these 35 ROIs overlap with those found via node persistence. Supplementary Table S3 lists ROIs with significant between-group differences identified by both node persistence and node frequency. For each of the 200 ROIs, group-wise averages of node persistence and node frequency are provided in Supplementary Table S4 along with FDR-corrected p-values, for both the MPI-LEMON and ABIDE-I datasets.

### Behavioral and cognitive relevance of region level differences

In the preceding subsection, we found that node persistence can identify 108 ROIs with significant differences between young and elderly individuals in the MPI-LEMON dataset, and 27 ROIs with significant differences between ASD and TD groups within the ABIDE-I dataset. These ROIs are distributed across multiple RSNs defined by the Schaefer atlas. Subsequently, to identify the cognitive domains related to these significant ROIs, we performed a Neurosynth meta-analysis (see Methods section) for each RSN. Only the ROIs exhibiting significant between-group differences in node persistence were considered in this analysis.

In the MPI-LEMON dataset, our analysis primarily focused on somatomotor, dorsal attention, salience/ventral attention, and default networks, as these RSNs not only exhibit significant differences across three global measures (persistent entropy, *L*^1^-norm, and *L*^2^-norm) but also contain the majority of ROIs with significant differences in node persistence. For the ABIDE-I dataset, we focused on the somatomotor and default networks, as they exhibit significant group differences at the RSN-level and also contain the majority of ROIs with significant differences identified via node persistence. Although the salience/ventral attention network also showed significant group differences at the RSN-level in ABIDE-I, it is excluded here as only one region is identified using node persistence.

The word clouds in Fig. 5(a) illustrate the behavioral relevance of the key brain regions corresponding to the four RSNs for the MPI-LEMON dataset. We found that ROIs with age-related changes in node persistence are associated with movement in the somatomotor and dorsal attention networks, somatosensory and affective processing in the salience/ventral attention network, and language, social cognition, and memory in the default network. The word clouds in Fig. 5(b) illustrate the behavioral relevance of the key brain regions corresponding to the somatomotor and default networks for the ABIDE-I dataset. We found that ROIs with ASD-related differences in node persistence are associated with movement in the somatomotor network and social cognition in the default network. The terms linked to significantly different ROIs across all seven RSNs for both datasets are summarized in Supplementary Table S5.

**FIG. 5.**
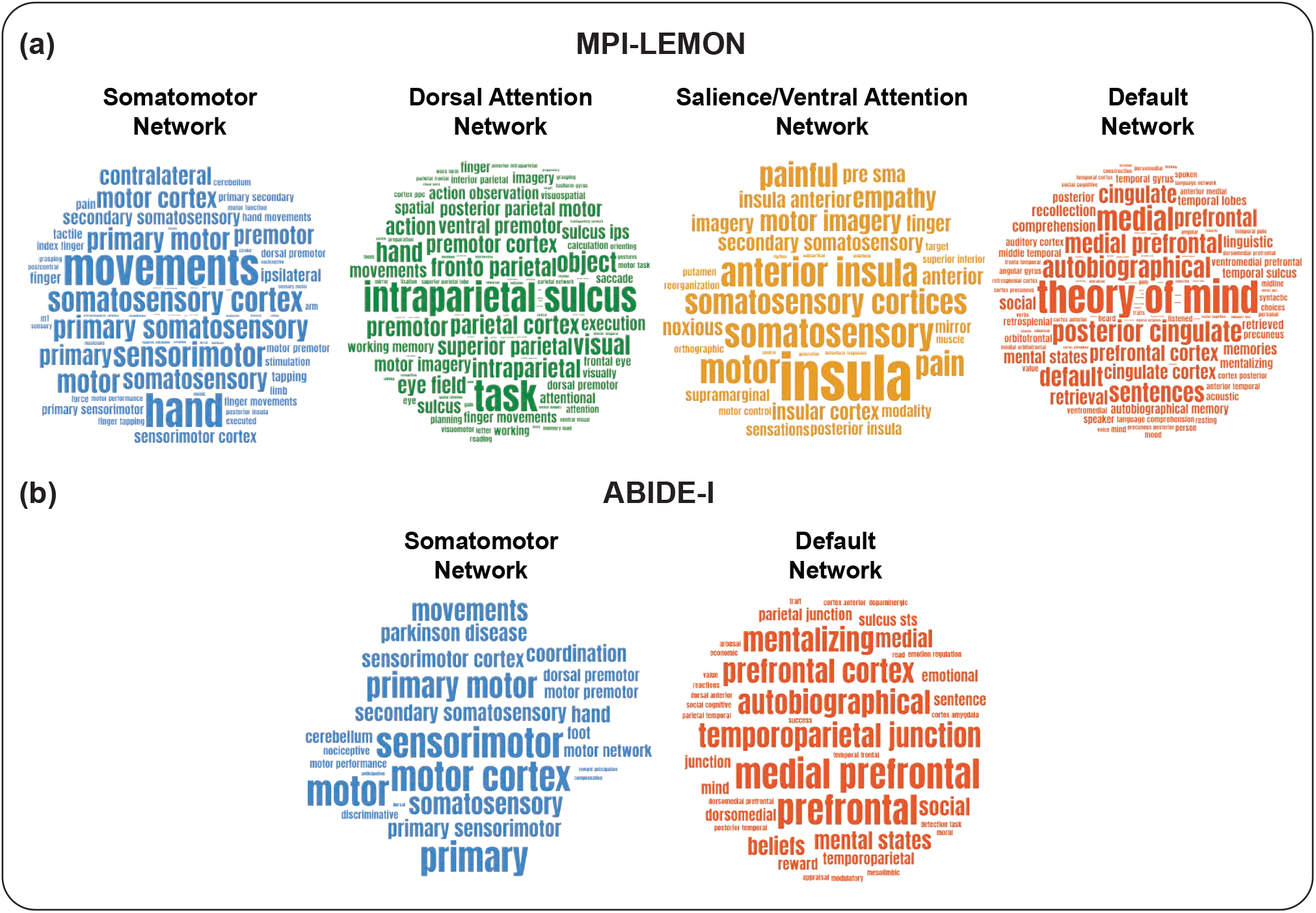
Behavioral and cognitive relevance of ROI-level changes using Neurosynth meta-analysis decoding. (a) MPI-LEMON dataset: word clouds highlighting cognitive and behavioral terms related to significantly different age-related brain regions identified via node persistence in four RSNs: somatomotor, dorsal attention, salience/ventral attention, and default network. (b) ABIDE-I dataset: word clouds highlighting cognitive and behavioral terms related to significantly different brain regions (*p <* 0.05, FDR-corrected) between individuals with autism spectrum disorder (ASD) and typically developing (TD) individuals as identified via node persistence, in two RSNs: somatomotor and default network. The words are scaled according to their frequency of occurrence. However, the scaling is performed independently for each word cloud. In other words, frequencies are not visually comparable across different word clouds. The word clouds are created using the online tool https://www.wordclouds.com.

### Correlation of topological measures with phenotypic and clinical scores

We conducted a correlation analysis to examine the relationship of PH-based metrics with phenotypic test scores for the MPI-LEMON dataset and clinical scores of symptom severity for individuals with ASD in the ABIDE-I dataset. The correlation analysis was performed for topological measures at all three scales.

MPI-LEMON dataset provides 52 affective processing related scores corresponding to all 225 subjects. These scores are divided across eight tests: (a) Cognitive Emotion Regulation Questionnaire (CERQ, 9 phenotypic scores), (b) Coping Orientations to Problems Experienced (COPE, 14 phenotypic scores), (c) Emotion Regulation Questionnaire (ERQ, 2 phenotypic scores), (d) Measure of Affect Regulation Style (MARS, 6 phenotypic scores), (e) Perceived Stress Questionnaire (PSQ, 5 phenotypic scores), (f) State-Trait-Angstinventar (STAI-G-X2, 1 phenotypic score), (g) Trait Emotional Intelligence Questionnaire-Short Form (TEIQue-SF, 5 phenotypic scores), and (h) Trierer Inventar zum Chronischen Stress (TICS, 10 phenotypic scores).

In the MPI-LEMON dataset, we found that PH-based measures exhibit significant correlations (*p* < 0.05, FDR-corrected) with TICS scores at the global level as well as at the level of RSNs. At the global level, persistent entropy shows significant correlations with 6 out of 10 phenotypic TICS scores, while the *L*^1^-norm and *L*^2^-norm show significant correlations with 1 and 2 scores, respectively. At the RSN-level, these measures show significant correlations with TICS scores within the visual, dorsal attention, and salience/ventral attention networks. Specifically, in the visual network, persistent entropy shows significant correlations with 1 out of 10 TICS scores, whereas both *L*^1^-norm and *L*^2^-norm exhibit significant correlations with 9 out of 10 TICS scores. In the dorsal attention network, persistent entropy and the *L*^1^-norm show significant correlations with 1 and 3 out of 10 TICS scores, respectively. In the salience/ventral attention network, persistent entropy is significantly correlated with 5 TICS scores, while the *L*^1^-norm and *L*^2^-norm exhibit significant correlations with 1 and 2 TICS scores, respectively. Supplementary Table S6 lists all the correlations between 52 affective processing related scores and three global measures at both the global and RSN levels.

At the ROI-level, age-related differences detected via node persistence in the salience/ventral attention network are related to affective and somatosensory processing (see Fig. 5(a)). Next, we examined the ROI-level correlations by considering the ROIs with significant between-group differences identified via node persistence. In the salience/ventral attention network, 11 such regions were present, which leads to 11 × 10 = 110 correlations. Of these, 21 exhibit significant correlations (*p* < 0.05) before FDR correction. These 21 significant correlations are distributed among seven brain regions. After FDR correction, two ROIs LH SalVentAttn FrOperIns 1 and RH SalVentAttn FrOperIns 3 show significant correlation with node persistence. All the significant correlations are positive, indicating that higher values of PH-based measures are associated with higher levels of chronic stress. Supplementary Table S7 provides the ROIs that exhibit significant correlations between all the test scores and node persistence.

In the ABIDE-I dataset, we identified two clinical scores based on the Autism Diagnostic Interview-Revised (ADI-R) criteria that assess symptom severity in individuals with ASD: (a) verbal and (b) social scores. Our analysis revealed no significant correlations between topology-based measures and clinical scores at the global or RSN-level. Correlations between the two clinical scores and the three PH-based measures at both global and RSN-level are provided in Supplementary Table S8. We further explored ROI-level correlations by focusing on ROIs that showed significant between-group differences based on node persistence. No significant correlations were observed between clinical scores and node persistence at the ROI-level (see Supplementary Table S7).

### Linking ROI-level differences in topological measures with non-invasive brain stimulation outcomes

Alongside the meta-analysis decoding, we conducted an additional analysis to evaluate the relevance of our findings with the existing literature on non-invasive brain stimulation (NIBS) in healthy elderly individuals and individuals with ASD. We focused on three commonly used NIBS techniques, namely transcranial direct current stimulation (tDCS), transcranial alternating current stimulation (tACS), and transcranial magnetic stimulation (TMS). For the MPI-LEMON dataset, we investigated the overlap between brain regions exhibiting differences in PH-based measures and those where NIBS has been shown to enhance motor performance in healthy elderly individuals. For the ABIDE-I dataset, we investigated the overlap with brain regions where NIBS has indicated reductions in ASD-related symptoms. The sets of brain regions where NIBS interventions have shown beneficial outcomes were curated in two prior studies conducted by some of us [67, 69].

The data derived from previous NIBS studies in healthy elderly individuals reveal four cortical target regions with evidence for improvement in motor function, namely primary motor cortex, dorsolateral prefrontal cortex, posterior parietal cortex, and right supplementary motor area. In the Schaefer atlas, these target regions correspond to a total of 42 ROIs. These ROIs are distributed across five RSNs: somatomotor (11), dorsal attention (12), salience/ventral attention (4), control (8), and default (7) networks. The Euler diagram in Fig. 6(a) illustrates the overlaps among 42 ROIs exhibiting improvements in motor function following NIBS, along with the ROIs identified by node persistence and node frequency. Additionally, it highlights both the overlaps and unique ROIs identified across NIBS studies, node persistence, and node frequency. Among these 42 clinically relevant ROIs, 27 are detected using node persistence and 10 are detected using node frequency. The clinically relevant ROIs detected by node persistence and node frequency are distributed across five and four RSNs, respectively. No clinically significant ROIs are found within the visual and limbic networks (see Supplementary Table S9). Further, the 10 clinically relevant ROIs detected by node frequency are a subset of those detected by node persistence (see Fig. 6(a) and Supplementary Table S9). In other words, node persistence uniquely identified 17 clinically relevant ROIs.

**FIG. 6.**
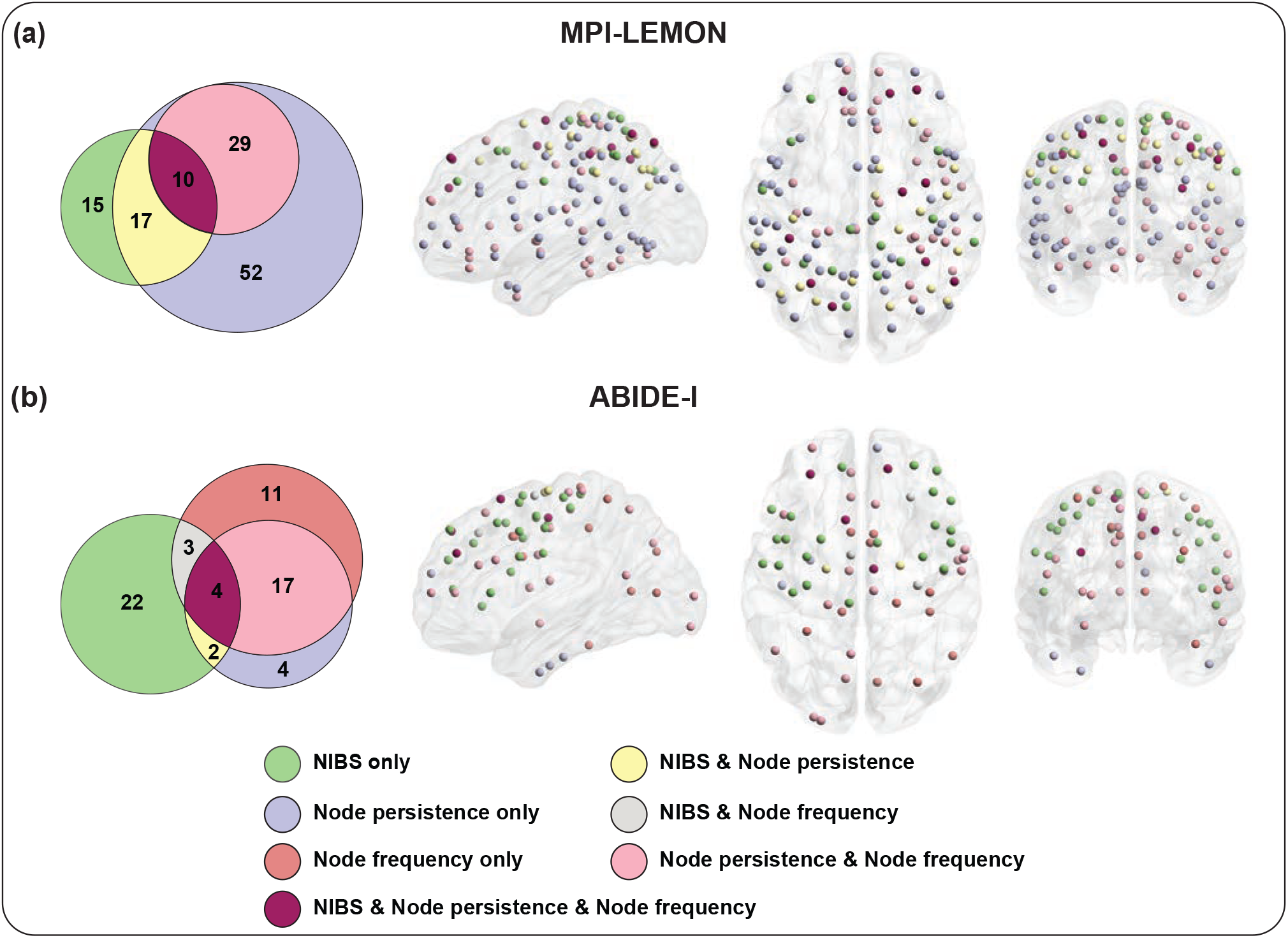
Visual representation of brain regions identified via non-invasive brain stimulation (NIBS) and local persistent homology-based measures. (a) MPI-LEMON dataset: Euler diagram depicting overlaps between brain regions identified by NIBS (42), node persistence (108), and node frequency (39), along with unique regions in each set. Accompanying this, the brain regions corresponding to each partition of the Euler diagram are visually represented. Within NIBS, 27 of 42 ROIs are identified considering both node persistence and node frequency. 17 ROIs are uniquely identified using node persistence. 10 ROIs identified via node frequency form a subset of ROIs detected by node persistence. (b) ABIDE-I dataset: Euler diagram depicting overlaps between brain regions identified by NIBS (31), node persistence (27), and node frequency (35), along with unique regions in each set. Accompanying this, the brain regions corresponding to each partition of the Euler diagram are visually represented. Within the 31 ROIs in NIBS, 9 ROIs are identified considering both node persistence and node frequency. Two and three ROIs are uniquely identified considering node persistence and node frequency, respectively.

The data derived from previous NIBS studies in individuals with ASD revealed five cortical target regions that show evidence for improving behavioral or cognitive symptoms associated with ASD, namely premotor cortex, DLPFC, pars triangularis, pars opercularis, and left primary motor cortex. These target regions encompass 31 Schaefer ROIs, and are distributed across five RSNs: somatomotor (10), dorsal attention (5), salience/ventral attention (4), control (5), and default (7) networks. The Euler diagram in Fig. 6(b) shows the overlaps among the 31 ROIs exhibiting ASD related improvements following NIBS and ROIs identified by PH-based local measures. It also illustrates both the overlaps and unique ROIs identified across NIBS studies, node persistence, and node frequency. Among these 31 clinically relevant ROIs, 6 are detected using node persistence and 7 are detected considering node frequency. These clinically relevant ROIs are distributed across three RSNs: somatomotor, salience/ventral attention, and default networks. Both the local measures collectively identified 9 out of 31 clinically relevant ROIs, with 2 uniquely identified by node persistence and 3 uniquely detected by node frequency (see Fig. 6(b) and Supplementary Table S9).

### Comparison with existing local measures based on persistent homology

Petri *et al*. [49] introduced the concept of homological scaffolds, which are secondary networks that summarize one-dimensional holes (cycles) captured by PH. Building on this idea, nodal Persistence Scaffold Strength (nodal PSS) was proposed as a node-level centrality measure that quantifies the contribution of a region to these cycles [50].

Specifically, nodal PSS considers the edges involved in representative *H*_1_ cycles extracted from PH via JavaPlex [79], and assigns strength scores to nodes accordingly. Other local topological measures, such as local PH, can also be defined at the level of individual nodes in a simplicial complex [64]. In this study, we included nodal PSS as a reference method to compare with our newly proposed local topological measures of node persistence and node frequency, since all these measures characterize the structure of one-dimensional holes. Further, nodal PSS has also been employed in the context of fMRI data analysis [50].

Notably, both node persistence and node frequency are less computationally expensive than the construction of homological scaffolds. The latter is significantly more time-consuming, particularly during the filtration process, where building the simplicial complex becomes increasingly intensive for larger point sets. This makes node persistence and node frequency better suited for node-level PH analyses on larger simplicial complexes compared to nodal PSS. In particular, when applied to FC matrices with 200 regions, we found that the scalability of homological scaffold construction significantly diminishes. Therefore, to identify ROI-level changes in functional connectivity using nodal PSS, we computed nodal PSS by considering FC matrices at the level of RSNs. Specifically, the homological scaffolds were constructed independently for each RSN, and nodal PSS was computed for each ROI within these networks. To ensure a consistent comparison with our proposed measures, we also recomputed node persistence and node frequency using FC matrices restricted to individual RSNs. In other words, our comparative analysis focused solely on intra-RSN topological patterns by excluding inter-RSN connections, thereby enabling the detection of ROI-level differences within each RSN. A two-tailed two-sample t-test was utilized to detect statistical differences between the groups, and FDR correction was applied independently for each RSN.

In the MPI-LEMON dataset, node persistence, node frequency, and nodal PSS identify age-related differences in 59, 40, and 58 ROIs, respectively (*p <* 0.05, FDR-corrected). In the ABIDE-I dataset, these three measures reveal significant ASD-related differences in 53, 35, and 45 ROIs, respectively (*p <* 0.05, FDR-corrected). The overlaps among these sets and their distribution across RSNs are provided in the Supplementary Text. All significant ROIs identified using node persistence, node frequency, and nodal PSS for both datasets are listed in Supplementary Table S10. Additionally, Supplementary Table S11 provides the group-wise averages of node persistence, node frequency, and nodal PSS for each of the 200 ROIs across both datasets, along with the corresponding FDR-corrected *p*-values.

Finally, we investigated the overlap between our RSN-to-local level analysis results and clinically validated ROIs from the NIBS literature. The UpSet plots [80] in Supplementary Fig. S6 illustrate the number of ROIs associated with clinical improvement and how many of these were identified using node persistence, node frequency, and nodal PSS, along with their intersections, for both MPI-LEMON and ABIDE-I datasets. In the MPI-LEMON dataset, of the 42 clinically relevant regions identified through NIBS, node persistence, node frequency, and nodal PSS detected 15, 13, and 18 ROIs, respectively. Similarly, in the ABIDE-I dataset, out of the 31 clinically relevant ROIs identified by NIBS, the three measures identified 9, 3, and 7 ROIs, respectively. Further details on the overlap between regions identified by local PH measures and NIBS-targeted regions are provided in the Supplementary Text and Supplementary Table S9.

To summarize, when examining ROI-level differences in functional connectivity during healthy aging and ASD using RSN-level FC matrices, node persistence identifies more brain regions than nodal PSS. In determining clinically relevant regions, nodal PSS detects more regions than node persistence in healthy aging, whereas in autism, node persistence identifies more regions than nodal PSS. However, both node persistence and nodal PSS can uniquely identify certain clinically relevant ROIs across both datasets.

## DISCUSSION AND CONCLUSION

To the best of our knowledge, this study is the first attempt to apply PH, a key tool in TDA, to analyze brain functional connectivity in individuals undergoing healthy aging and those with ASD at three different scales: (a) global (brain-wide changes), (b) mesoscopic (RSN-level changes), and (c) local (ROI-level changes). In addition, we introduce a scalable and computationally efficient method for extracting local topological insights from brain functional connectivity using PH. At the global level, our analysis using PH-based measures revealed clear distinctions between healthy young and elderly individuals, as well as between ASD and TD individuals. Our mesoscale comparisons revealed specific RSNs that are more sensitive to age or ASD related changes. Finally, our proposed local metrics based on PH successfully identified brain regions associated with healthy aging and ASD, highlighting their potential to uncover local topological changes in functional connectivity.

In this study, we acquired FC matrices derived from the MPI-LEMON and ABIDE-I datasets to investigate changes in brain functional connectivity associated with healthy aging and autism, respectively. The FC matrix, representing pairwise Pearson correlations among 200 brain regions as specified by the Schaefer atlas, served as the initial input for our PH-based analysis. We focused on functionally meaningful interactions by retaining only positive correlations, which were subsequently transformed into ultrametric distance matrices. This transformation allowed us to compute PH-based measures via Rips filtration. In particular, we employed persistent entropy of the persistence barcodes, *L*^1^-norm and *L*^2^-norm of the persistence landscape, for both brain-wide and RSN-level investigations. For the ROI-level analysis, we employed our proposed metrics, node persistence and node frequency, to identify specific ROIs responsible for functional connectivity differences. Remarkably, our methodology enabled the extraction of multiscale topological features from FC matrices, revealing both the intrinsic RSNs as well as specific brain regions responsible for alterations in resting-state functional connectivity associated with healthy aging and ASD.

At the global scale, our analysis demonstrated significant differences in topological measures between the groups utilizing 200 *×* 200 FC matrices. In the context of healthy aging, the young group exhibited higher persistent entropy, as well as higher *L*^1^-norm and *L*^2^-norm values compared to the elderly group. Further, individuals with ASD showed increased persistent entropy but reduced *L*^1^-norm and *L*^2^-norm values relative to the TD group. A higher persistent entropy indicates a more uniform distribution of the persistence of features across the filtration, suggesting the absence of dominant topological structures. Conversely, a lower persistent entropy reflects a concentration of longer-lived features, implying the presence of prominent topological structures within the data [75]. Higher *L*^1^-norm and *L*^2^-norms of the persistence landscape indicate that one-dimensional topological features persist longer, capturing their dominance and longevity across the filtration. Additionally, 1-Wasserstein, 2-Wasserstein, and bottleneck distances revealed higher inter-group distances compared to intra-group distances for both the MPI-LEMON and ABIDE-I datasets, indicating a greater topological similarity within groups than between groups.

At the mesoscopic scale, we analyzed each of the seven RSNs individually, focusing only on the intra-RSN functional connectivity, specifically: visual, somatomotor, dorsal attention, salience/ventral attention, limbic, control, and default networks. In the MPI-LEMON dataset, RSN-level comparison of persistent entropy, *L*^1^-norm, and *L*^2^-norm revealed that healthy aging predominantly alters the topology of four RSNs: somatomotor, dorsal attention, salience/ventral attention, and default networks. In the ABIDE-I dataset, these global measures revealed that ASD primarily affects the topology of three RSNs: somatomotor, salience/ventral attention, and default networks. Therefore, our mesoscale analysis revealed that, although significant topological differences were observed at the global level, they do not arise uniformly across all RSNs. In other words, only specific RSNs contribute to the observed effects.

To identify region-level differences associated with healthy aging and ASD, our proposed metrics, node persistence and node frequency, were employed. Using node persistence, we identified 108 and 27 brain regions with significant age-related and ASD-related changes, respectively. In the MPI-LEMON dataset, brain regions with age-related changes were concentrated across four RSNs: somatomotor, dorsal attention, salience/ventral attention, and default networks. In the ABIDE-I dataset, brain regions with ASD-related changes were concentrated across the somatomotor and default networks. Remarkably, these RSNs also exhibited significant group differences during our mesoscale analysis, which highlights the consistency of our findings across different scales. On the other hand, node frequency identified 39 and 35 brain regions with significant age-related and ASD-related changes, respectively. Notably, in the healthy aging study, all 39 brain regions identified by node frequency were also captured by node persistence, indicating strong agreement between the two measures. In contrast, in the ASD study, only 21 out of the 35 ROIs identified by node frequency overlapped with those identified by node persistence, indicating that the two measures may capture distinct aspects of region-level topological differences in ASD.

Next, we used Neurosynth meta-analysis decoding to determine the behavioral and cognitive relevance of the changes arising due to healthy aging and ASD, as identified by node persistence. Our analysis revealed that the brain regions exhibiting age-related differences in node persistence are mainly related to movement, language, social cognition, memory, somatosensory, and affective processing. Previous research suggests that aging impacts various domains of motor performance [81, 82], including coordination [83], movement variability [84], and speed [85], as well as language production [86] and affective processing [87]. It also leads to diminished somatosensory functions such as sensitivity to warmth, touch, and vibration [88], collectively highlighting the physical and emotional changes associated with aging [87]. Our analysis revealed that the brain regions exhibiting ASD-related differences in node persistence are mainly related to movement and social cognition. Previous meta-analysis decoding suggests that ASD influences the cognitive domains [16–18, 89–95]. Hence, our findings indicate that regions exhibiting differences in node persistence in both healthy aging and ASD are associated with cognitive domains and abilities that are commonly reported to undergo age-related or ASD-related changes.

Following the meta-analysis, we assessed the alignment between our findings and prior tDCS, tACS, and TMS studies. Our analysis revealed that brain regions exhibiting altered PH-based local measures in both healthy elderly individuals and those with ASD overlap with regions previously shown to benefit from non-invasive stimulation, either through improved motor performance in aging or reduction in symptom severity in ASD. Furthermore, in determining clinically significant brain regions, node persistence proved superior in detecting a larger number of clinically relevant brain regions in the context of healthy aging, while for autism, node persistence and node frequency revealed a comparable number of clinically relevant brain regions. To the best of our knowledge, this study makes the first attempt to validate PH-based analyses of brain functional connectivity networks using evidence from non-invasive brain stimulation (tDCS/tACS/TMS) studies. Our findings suggest that node persistence effectively captures atypical connectivity patterns in clinically meaningful brain regions across both healthy aging and ASD. More broadly, our results support the potential of PH-based measures to generate hypotheses about clinically relevant targets, which can subsequently be evaluated through non-invasive stimulation protocols.

We compared our proposed local topological measures with nodal PSS, an existing method that also characterizes one-dimensional holes in functional connectivity networks. Our results show that node persistence consistently identifies a larger number of brain regions with significant group differences compared to nodal PSS in both the MPI-LEMON and ABIDE-I datasets. However, when aligning our findings with non-invasive stimulation outcomes, we found that the regions highlighted by node persistence do not entirely overlap with those identified by nodal PSS. This suggests that different local topological measures may offer complementary insights into clinically relevant brain regions. Due to the higher computational complexity of nodal PSS, these comparisons were conducted using FC matrices restricted to individual RSNs. In contrast, our proposed measures, node persistence and node frequency, are scalable and were applied to FC matrices involving all ROIs in the main analyses reported in this study. This scalability and methodological flexibility make our approach well-suited for neuroimaging studies involving large populations.

In conclusion, we found that PH facilitates the identification of age-related as well as ASD-related changes in functional connectivity at multiple spatial scales. Further, we introduced two local PH-based measures, namely node persistence and node frequency, to detect topology-influenced variations at the level of individual brain regions. Moreover, to reinforce the clinical relevance of our findings for non-invasive brain stimulation, we demonstrated that local measures based on PH can effectively identify brain regions impacted by healthy aging and ASD. One limitation of our approach is that the node-level measures currently consider only one-dimensional topological features such as loops or cycles, potentially overlooking more complex structures. Future research could extend this framework to incorporate higher-dimensional features, such as voids, for a more comprehensive characterization of network topology. Furthermore, node persistence could be utilized to examine local characteristics of brain functional connectivity networks beyond aging and ASD, and could potentially be extended to other types of networks, such as task-based fMRI networks.

## METHODS

In this study, we utilize PH-based metrics to investigate changes in resting-state functional connectivity in individuals undergoing healthy aging and those with autism spectrum disorder (ASD). First, we acquired functional connectivity (FC) matrices obtained from resting-state functional MRI (rs-fMRI) images of subjects from the MPI-LEMON [66] and ABIDE-I [68]. Second, we constructed a sequence of nested Vietoris-Rips (Rips) complexes [96] on the distance matrix derived from each FC matrix. Third, we computed topological invariants based on PH to identify the changes in functional connectivity at three different scales: (a) global scale (brain-wide changes), (b) mesoscopic scale (RSN-level changes), and (c) local scale (ROI-level changes).

### Functional Connectivity Matrices

The functional connectivity (FC) matrices used in this study are obtained from preprocessed rs-fMRI images of subjects from the MPI-LEMON and ABIDE-I datasets. These FC matrices were previously generated by some of us using the CONN functional connectivity toolbox [97], and are available at https://github.com/asamallab/Curvature-FCN-Aging and https://github.com/asamallab/Curvature-FCN-ASD. The FC matrix of each subject is a 200 *×* 200 square matrix representing the pairwise correlations among the 200 regions of interest (ROIs) as outlined in the Schaefer atlas [72]. Each matrix entry (*i, j*) contains a numerical value representing the Pearson correlation between the time series of ROI *i* and ROI *j*. The time series for each ROI was computed by averaging the time series of all voxels located within that region from the preprocessed fMRI images. For a comprehensive overview of the preprocessing pipeline and FC matrix generation, readers are referred to [67, 69].

In addition to assigning each voxel to one of 200 ROIs, the Schaefer atlas [72] also associates each ROI with one of seven resting-state networks (RSNs). These RSNs include the *visual network* (29 ROIs), *somatomotor network* (35 ROIs), *dorsal attention network* (26 ROIs), *salience/ventral attention network* (22 ROIs), *limbic network* (12 ROIs), *control network* (30 ROIs), and *default network* (46 ROIs). This parcellation method allows for examining brain functional connectivity at the local scale (ROI-level) as well as at the mesoscopic scale (RSN-level).

A summary of the datasets obtained for this study is as follows:

- **MPI-LEMON**: The FC matrices of 225 participants from the Max Planck Institute Leipzig Study for Mind-Body-Emotion Interactions (MPI-LEMON) dataset [66] consists of 153 healthy young (age range 20− 35 years) and 72 healthy elderly individuals (age range: 59 *−* 77 years) [67]. Notably, MPI-LEMON dataset does not include any middle-aged individuals.
- **ABIDE-I**: The FC matrices of 820 participants (age range: 7 −64 years) from the ABIDE-I project [68], consists of 395 individuals with ASD and 425 age-matched TD individuals [69].

### Topological representation

#### Simplicial complex

A simplicial complex is a higher-dimensional generalization of a graph, constructed by combining vertices, edges, triangles, tetrahedra, and their higher-dimensional counterparts. Mathematically, a simplicial complex is defined as a collection *K* of non-empty subsets of a finite set *V* that satisfies the following two conditions:

1. If *σ* ∈ *K* and *τ* ⊊ *σ*, then *τ* must also be an element of *K*.
2. For each *v* ∈ *V*, the singleton {*v*} is included in *K*.

The elements of *V* are called the *vertices* of *K*, while the elements of *K* are referred to as *simplices*. If *τ* ⊊ *σ* for simplices *τ, σ* ∈ *K*, then *τ* is termed a *face* of *σ* and denoted as *τ < σ*. A simplex containing *p* + 1 elements is known as a *p-simplex*, meaning it has dimension *p*. The set of all *p*-simplices in *K* is denoted by *K*_*p*_. The *dimension* of the simplicial complex *K* is defined as the highest dimension among its simplices. To explore simplicial complexes in greater detail, we recommend standard texts [28, 98] in algebraic topology.

#### Vietoris-Rips complex

Let (*X, d*) be a metric space and *S* a finite set of points in the metric space, often referred to as a point cloud. For a given radius *ϵ* (*>* 0) the Vietoris–Rips complex or Rips complex *R*_*ϵ*_(*S*) is a simplicial complex made up of vertices from *S*, with simplices formed from finite subsets of *S*, where each pair of points in a given simplex has a distance at most 2*ϵ*. Formally, Rips complex *R*_*ϵ*_(*S*) is defined as,

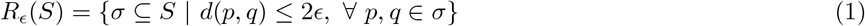

where *d*(*p, q*) denotes the distance between points *p* and *q*. A 1-simplex (or edge) between two points is added if the distance between them is ≤ 2*ϵ*. Given three points, a 2-simplex (or filled triangle) is formed if the pairwise distances between them are all less than or equal to 2*ϵ*. In the case of higher-dimensional simplices, a *k*-simplex arises from a subset of (*k* + 1) points in *S* if every pair of points in the subset is connected by an edge.

Rips complex is an efficient method that enables the construction of a simplicial complex on an arbitrary space. Besides Rips complex, there are other methods available, such as Čech complex and alpha complex to construct a simplicial complex [99]. However, Rips complex is more computationally efficient than Čech complex and is often regarded as an approximation of the latter [29, 100, 101]. The efficiency arises from the simplicity of its construction, as determining whether a simplex *σ* ⊆*S* belongs to *R*_*ϵ*_(*S*) requires only the computation of pairwise distances. For more details regarding Rips complex see e.g. [32, 96].

#### Construction of Rips complex from FC matrix

Given an FC matrix **C**, the *ultrametric distance matrix* [73] (**D**) is constructed using the distance measure given by Eq. (2).

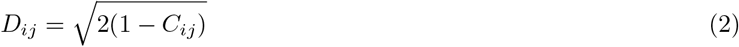

where *C*_*ij*_ is the Pearson correlation between the brain regions *i* and *j*. The ultrametric distance in our analyses is defined based solely on positive correlations in the FC matrix, reflecting the dominating role of positive connectivity in supporting overall brain function [74, 102]. As a result, each value in the distance matrix ranges between 0 and 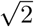. These distance matrices are utilized to construct the Rips complex [96] and to compute PH. Unique values from the distance matrix serve as a filtration parameter 2*ϵ* of the resulting sequence of Rips complexes.

### Persistent homology of simplicial complexes

#### Homology

The construction of homology groups of a simplicial complex *K* begins with a *p*-chain. A *p*-chain *C*_*p*_ for an oriented simplicial complex *K* is defined by the formal linear combinations of oriented *p*-simplices *α*_*i*_ of *K*, expressed as:

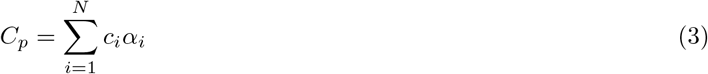

where the coefficients *c*_*i*_ are elements from a base field *F*. Under pointwise addition, the family of *p*-chains forms a group over the base field *F*, called the *p*-dimensional chain group of *K*, and this group is denoted by *C*_*p*_(*K*).

Next, the boundary operator *∂*_*p*_ : *C*_*p*_ → *C*_*p−*1_ maps the *p*-chain to the sum of the (*p*−1)-dimensional faces of that *p*-simplex resulting in a (*p* −1)-chain. Let *α* = [*x*_0_, *x*_1_, …, *x*_*p*_] be an oriented *p*-simplex, and let 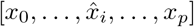 denote the (*p* − 1)-simplex after the removal of the point *x*_*i*_ from that *p*-simplex. Then the boundary operator *∂*_*p*_ is defined as follows:

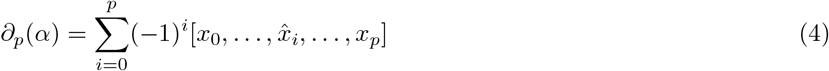

The boundary operator satisfies the fundamental property *∂*_*p*_ °*∂*_*p*+1_ = 0, which ensures that the boundary of a boundary vanishes. Now, sequentially arranging the chain groups and boundary operators yields a chain complex:

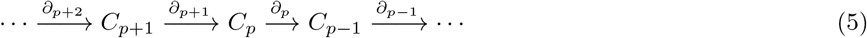

Next, *p*-cycles and *p*-boundaries are defined from the boundary operator *∂*_*p*_. The kernel of the boundary operator *∂*_*p*_ is called a *p*-cycle and is denoted by *Z*_*p*_, i.e.

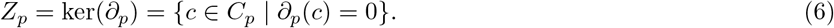

In other words, a *p*-cycle is a *p*-chain with an empty boundary. A *p*-boundary is a *p*-cycle that lies in the image of the boundary operator *∂*_*p*+1_. The *p*-boundaries form a group *B*_*p*_, which is a subgroup of the *p*-cycles *Z*_*p*_ [28], i.e.

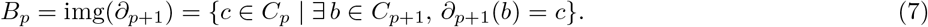

Thus, the simplicial *p*-homology group of *K* is defined by the quotient group,

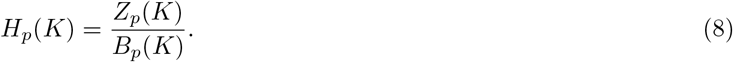

Since the coefficient field is commutative, *B*_*p*_ is a normal subgroup, and *H*_*p*_ is also a group.

The elements of the *p*-homology group *H*_*p*_ are informally known as *p*-holes. Notably, *H*_*p*_ forms a vector space over the field *F* and the dimension of *H*_*p*_ defines *β*_*p*_, the *p*-Betti number. Intuitively, this Betti number provides the number of holes formed by *p*-simplices. For instance, *β*_0_ represents the number of connected components, *β*_1_ represents the number of loops or one-dimensional holes, *β*_2_ represents the number of voids or cavities or two-dimensional holes, and this pattern continues for higher dimensions.

#### Filtration of simplicial complexes

As mentioned, a simplicial complex *K* is a set of finite number of simplices and the simplices are either disjoint or intersect in a common face. A subcomplex is a subset *L* ⊆ *K* of the set of simplices such that all the elements in *L* also satisfy the properties of a simplicial complex.

Given a simplicial complex *K*, a filtration of length *n* is a nested sequence of subcomplexes, where each subcomplex is contained within the next. Formally, it is defined as:

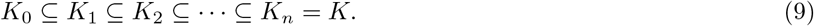

Here, *K*_*i*_ represents a simplicial complex at the *i*^th^ stage.

The Rips complex defined in the Methods section admits a natural filtration of simplicial complexes where each subcomplex is indexed by a real parameter *ϵ* (radius), corresponding to half the distance between pairs of points. Given that only a finite number of point pairs exist, there will be finitely many *ϵ* values at which new simplices get introduced. In the beginning of the filtration process, i.e. when *ϵ* is very small, the Rips complex contains only the set of points *K*_0_ in a given metric space. Next, the radius *ϵ* around each point is gradually increased, and at each step of the filtration, a simplicial complex is formed. As the radius *ϵ* increases, higher-dimensional simplices such as edges, triangles and tetrahedra form. For large *ϵ*, all points become interconnected, forming a single large simplex. In summary, a filtration tracks how topological features (such as connected components, loops, and voids) evolve across different scales.

#### Persistent homology

In the filtration of a simplicial complex *K*, each subcomplex *K*_*i*_ contains *p*-chains, *p*-cycles, *p*-boundaries, and a *p*-boundary operator acting on the *p*-chains. Therefore, the *j*-persistent *p*-homology of *K*_*i*_ is defined as,

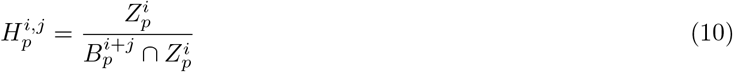

where 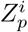 denotes the *p*-cycles in the subcomplex 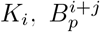 denotes the *p*-boundaries in *K*_*i*+*j*_. Therefore, the persistent *p*-Betti number of the subcomplex *K*_*i*_ becomes 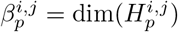.

The PH of a filtration of simplicial complexes gives more refined information than just the homology of the individual subcomplexes [26, 103, 104]. In particular, the PH group 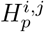 contains the *p*-cycles that form in the subcomplex *K*_*i*_ and become boundaries at the subcomplex *K*_*i*+*j*_. This indicates that the *p*-cycle persists from step *i* to step *i* + *j* as it appears (or is born) at *K*_*i*_ and disappears (or dies) at *K*_*i*+*j*_. In simple words, persistence captures the span of the important topological features from their appearance (birth) to their disappearance (death) across the filtration process. For comprehensive details on PH, refer to [26, 30, 31, 46, 99].

### Topological measures

#### Persistent entropy

PH measures the span of important topological features of a simplicial complex *K* such as the number of connected components (0-cycles), loops (1-cycles), voids (2-cycles), and so forth, throughout the filtration process. A visual representation of the span of these features can be provided in the form of *persistence barcodes*. Through persistence barcodes, complex high-dimensional data is distilled into a straightforward, interpretable format. In the filtration of a finite simplicial complex *K*, the *p*^th^ barcode diagram represents the birth and death of *p*-holes during the filtration process [29]. In particular, for a Vietoris-Rips filtration, a barcode spanning from *ϵ*_*i*_ to *ϵ*_*j*_ on the *x*-axis depicts a *p*-hole in *K*, with *ϵ*_*i*_ and *ϵ*_*j*_ representing its birth and death filtration values, respectively. The number of barcodes between *ϵ*_*i*_ and *ϵ*_*j*_ reflect the *p*-Betti number 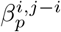.

*Persistent entropy* is a summary statistic of the persistence barcodes that quantifies the spans of the topological features from a sequence of birth–death information [75, 105, 106]. In particular, it provides the Shannon entropy of the persistence barcodes. Consider a persistence diagram *D* = {(*b*_*i*_, *d*_*i*_)}, where (*b*_*i*_, *d*_*i*_) represents the birth and death filtration values, with each *d*_*i*_ *<* +∞. Then the persistent entropy associated with *D* is defined as:

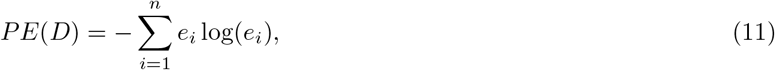

Where

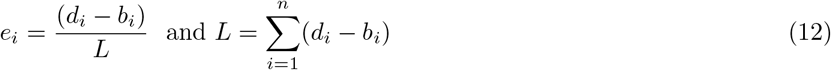

with *n* denoting the total number of bars and 0 ≤ *PE*(*D*) ≤ log(*n*). Persistent entropy shows how the spans of the features are spread out. A higher entropy value indicates that the spans of features are more evenly distributed.

#### Persistence diagram and persistence landscape

Like barcodes, a persistence diagram also provides a visual representation of the span of the features. A persistence diagram represents a multiset of points {(*b*_*i*_, *d*_*i*_)} in ℝ^2^, representing the birth and death pairs associated with *p*-holes [32]. In this diagram, each point (*b*_*i*_, *d*_*i*_) is plotted in the Cartesian plane, where the *x* and *y* axes represent filtration values at birth and death, respectively. Since *d*_*i*_ *> b*_*i*_, all points lie above the *b* = *d* line, and the farther above, the longer the span of the associated *p*-hole. Persistence diagrams are robust to noisy data [107]. Distance functions, such as the Wasserstein distance, can be considered to define a metric space of persistence diagrams [108, 109].

Next, to perform statistical analysis, a persistence diagram is transformed into a sequence of real-valued functions Λ_*i*_ : ℝ → ℝ for *i* ≥1, known as the persistence landscape [110], aggregating the essential information contained in persistence diagrams. The following method is used to construct a persistence landscape from the persistence diagram. First, a clockwise rotation of 45^*°*^ is applied to convert the diagonal of a persistence diagram into the *x*-axis. Then, isosceles right triangles are drawn from each point (*b*_*i*_, *d*_*i*_) of the persistence diagram, considering the point as the vertex. Next, a tent function is constructed from each of the generated triangles. For instance, the second tent function represents the second-highest value at each point across all triangles. In mathematical terms, a piece-wise linear function Λ_*i*_ : ℝ *→* [0, ∞) exists for every birth–death pair {(*b*_*i*_, *d*_*i*_)} in the persistence diagram defined as:

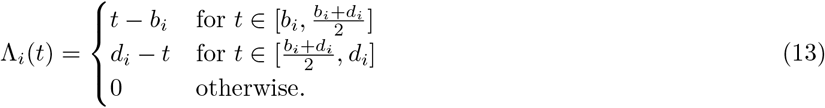

The *k*^th^ largest value among the set {Λ_*i*_} defines the persistence landscape *λ*_*k*_(*t*). In this study, we consider *k* = 1, as *λ*_1_(*t*) typically captures the most prominent topological features for each *t*. As a subset of a Banach space, the persistence landscape enables the calculation of *L*^*p*^-norms (1 ≤ *p* ≤ ∞), which is not the case for persistence diagrams. The *L*^*p*^-norms of a persistence landscape *λ*_1_(*t*) are defined as [76]:

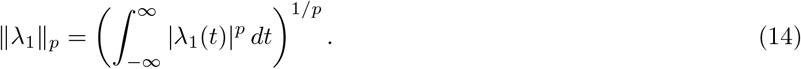

In the case of a single birth-death pair (*b, d*) in the persistence diagram, the norms of *λ*_1_(*t*) are given by: 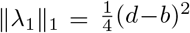 and 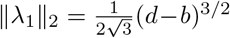. For multiple non-overlapping birth-death pairs {(*b*_*i*_, *d*_*i*_)}, the norms are computed as: 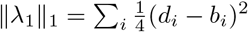 and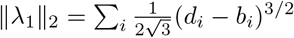.

Moreover, we calculate the Wasserstein and bottleneck distances to measure the difference between two persistence diagrams [32]. Let *X* and *Y* denote two persistence diagrams. The *p*-Wasserstein distance is the minimum total cost required to align points (birth-death pairs) from one diagram with another and is mathematically expressed as:

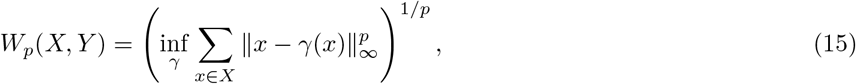

where *γ* represents a bijection between the points of *X* and *Y*, allowing points to be matched either to each other or, when necessary, to the diagonal, due to the difference in the cardinalities of *X* and *Y*. This distance quantifies disparities in topological features between two datasets. The bottleneck distance is a special case of the Wasserstein distance with parameter *p* → ∞, defined as:

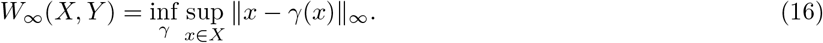

#### Node persistence and node frequency

Although PH captures multiscale topological information from datasets, it remains a global measure and, as such, is limited in its ability to characterize the topology of individual components. In the context of brain connectivity analysis, topological measures derived from PH, such as persistent entropy and persistent landscapes, may effectively distinguish global patterns in brain functional connectivity. However, they lack the ability to identify specific brain regions responsible for altered connectivity. In this section, we present two novel approaches to extract node-level information from the first homology group _1_, namely *node persistence* and *node frequency*.

A one-dimensional hole is defined by its one-dimensional boundary, which is a collection of 1-simplices (or edges) that form a cycle. However, there is no unique way to define the boundary of a hole. In other words, multiple cycles in the simplicial complex might describe the same one-dimensional hole [30]. As a result, it is crucial to consider a representative cycle for each class. In our work, we use the representative cycles defined by JavaPlex, an open-source library designed for computational topology and TDA, which is integrated into our workflow via Jython interface [79].

Let *G* = {*g*_1_, *g*_2_, …, *g*_*β*_} be the set of all one-dimensional holes, where the *i*^th^ one-dimensional hole is associated with the birth-death pair (*b*_*i*_, *d*_*i*_). The representative cycle of *g*_*i*_ is defined by the set of nodes. 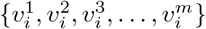 Furthermore, we use the term *persistence extent* to characterize the overall spread or range of persistence barcodes.

The *node persistence* of a node *v* is defined by aggregating the spans of all cycles that include the node over the persistence extent of the one-dimensional holes. Mathematically, it can be expressed as:

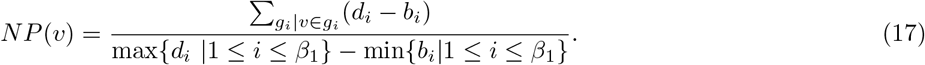

The *node frequency* of a node *v* provides the number of distinct cycles containing the node. Mathematically, it can be expressed as:

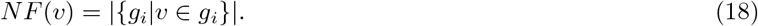

The two measures, node persistence and node frequency, yield node-level or local insights into PH while taking the one-dimensional holes into account.

### Persistent homology-based analysis of brain functional connectivity

We consider FC matrices from two datasets to conduct a PH-based analysis of brain connectivity, namely MPI-LEMON comprising 225 subjects, and ABIDE-I comprising 820 subjects. In particular, we employed topological measures derived from PH to compare resting-state functional connectivity between young and elderly groups in the MPI-LEMON dataset, and between individuals with ASD and TD controls in the ABIDE-I dataset.

#### Brain-wide analysis

To identify brain-wide or global changes in resting-state functional connectivity, we utilized the 200 *×*200 FC matrix for each subject. Each FC matrix was transformed into a distance matrix according to the procedure outlined in the Methods section. Subsequently, we constructed a filtration of Rips complexes over each distance matrix, indexed by the set of unique pairwise distances serving as the filtration parameters 2*ϵ*. Our analysis focused on the evolution of topological features as 2*ϵ* varies, captured by PH. These features include the zeroth homology group *H*_0_, which captures connected components, the first homology group *H*_1_, which corresponds to loops or one-dimensional holes, and the second homology group *H*_2_, which corresponds to voids or two-dimensional holes in the Rips complex [32, 96]. Particular attention was given to the *H*_1_ homology group. We calculated six global topological measures: (i) persistent entropy from the persistence barcodes associated with *H*_0_, *H*_1_, and *H*_2_ [75]; (ii) *L*^1^-norm of the persistence landscape associated with *H*_1_ [110]; (iii) *L*^2^-norm of the persistence landscape associated with *H*_1_; (iv) the 1-Wasserstein distance; (v) 2-Wasserstein distance; and (vi) bottleneck distance between the persistence diagrams associated with *H*_1_. We utilized the open source library Gudhi [111] to compute these global topological measures.

#### RSN-level analysis

As mentioned previously, the Schaefer atlas [72] consists of 200 ROIs or nodes which are partitioned across seven RSNs. To identify changes in resting-state functional connectivity at the RSN or mesoscale level, we focused on submatrices of the FC matrix containing entries corresponding to nodes within specific RSNs. Each submatrix was transformed into a distance matrix according to the procedure outlined in the Methods section. Consequently, for each subject, seven distinct distance matrices were constructed, each corresponding to one of the seven RSNs. This led to a total of 225 × 7 = 1575 and 820 × 7 = 5740 distance matrices for the MPI-LEMON and ABIDE-I datasets, respectively. The size of each distance matrix varies based on the number of nodes present in the respective RSN. Finally, we constructed a filtration of Rips complexes for each RSN over the corresponding distance matrix. Note that the construction of Rips complex for the RSN-level analysis follows the same approach as that employed in brain-wide analysis. However, in this scenario, only intra-RSN connections are taken into account. The six topological measures listed in brain-wide analysis were also calculated for determining RSN-level changes in functional connectivity. These include persistent entropy, *L*^1^-norm, *L*^2^-norm, 1-Wasserstein, 2-Wasserstein, and bottleneck distances.

#### ROI-level analysis

To identify ROI-level or local changes in resting-state functional connectivity, we utilized the filtration of Rips complexes generated during the brain-wide analysis. Subsequently, we calculated three topological measures based on PH: (i) node persistence, (ii) node frequency, and (iii) nodal PSS. These measures characterize node-level topological information by extracting the representative cycles of the first homology group *H*_1_, which corresponds to one-dimensional holes or cycles within the Rips complex. Node persistence and node frequency were computed via JavaPlex [79], while nodal PSS was computed using the Python package Holes [49, 112].

### Neurosynth meta-analysis decoding

Using the ROI-level analysis described in the preceding section, we identified specific brain regions associated with altered resting-state functional connectivity in healthy aging (based on the MPI-LEMON dataset) as well as during ASD (based on the ABIDE-I dataset). The cognitive and behavioral implications of these findings were then examined through the Neurosynth meta-analysis decoding [70, 71]. First, we used the Neurosynth meta-analysis tool to identify the terms that are related to behavior, cognition, and perception based on the centroid coordinates of the 200 ROIs in the Schaefer atlas. Second, we focused on nodes exhibiting significant inter-group differences in node persistence values. Each ROI or node belongs to one of the seven RSNs, as specified by the Schaefer atlas. Third, we computed the frequency of terms associated with significant nodes within each RSN. The statistical significance of these frequency counts was assessed using a procedure described in the following section.

### Statistical analyses

We performed two-tailed two-sample t-tests [77] to examine the statistical significance of the differences in the means of the measured values between (i) young and elderly groups in the MPI-LEMON dataset, and (ii) ASD and TD groups in the ABIDE-I dataset. To evaluate group-level differences at global and mesoscopic scales, comparisons were performed across three measures: persistent entropy, *L*^1^-norm, and *L*^2^-norm. The 1-Wasserstein, 2-Wasserstein, and bottleneck distance distributions are also compared between intra-group and inter-group conditions to assess whether persistence diagrams are more similar within groups (intra-group) than between them (inter-group). However, for these measures we performed one-tailed two-sample t-tests, as here our null hypothesis is that intra-group distances will be less than the inter-group distances. Furthermore, to evaluate group-level differences at the local scale, two-tailed two-sample t-tests are performed across all 200 nodes, for each of the three node-based measures: node persistence, node frequency, and nodal PSS.

To assess the statistical significance of the frequency of terms in the Neurosynth meta-analysis decoding, we compared the occurrences of the same terms within an equal number of randomly chosen nodes (proxy ROIs). For instance, considering node persistence, we obtained 108 nodes that exhibit significant differences between the young and elderly groups in the MPI-LEMON dataset. Therefore, from the 200 nodes in the Schaefer atlas, we randomly considered 108 nodes. This allowed us to establish a null distribution for term occurrences by generating 1000 sets of these randomly selected nodes. We then determined the z-score for each term’s frequency in the original set of ROIs. Thereafter, we convert these z-scores into p-values assuming a normal distribution [67, 69, 71].

To account for multiple comparisons and reduce the likelihood of false positives, we implemented a False Discovery Rate (FDR) correction [113] to adjust the p-values. We conducted all statistical tests using the SciPy [114] and statsmodels [115] packages in Python. Moreover, the analyses are performed without incorporating any covariates such as gender or age.

### Correlation of topological measures with cognitive, behavioral, and clinical scores

A correlation analysis was conducted to quantify the strength of the linear association between PH-based measures and (i) performance on cognitive/behavioral tests in the MPI-LEMON dataset, and (ii) clinical scores indicating symptom severity in the ABIDE-I dataset. We examined these correlations across all three spatial scales: (a) global scale (brain-wide changes), (b) mesoscopic scale (RSN-level changes), and (c) local scale (ROI-level changes).

The MPI-LEMON dataset includes comprehensive data on 6 cognitive tests and 21 questionnaires addressing emotional tendencies, personality traits, dietary habits, and addiction, accessible at https://fcon_1000.projects.nitrc.org/indi/retro/MPI_LEMON.html. We retrieved the phenotypic test scores corresponding to all 225 individuals. Subsequently, we calculated Spearman correlation of metrics derived from PH: persistent entropy, *L*^1^-norm, and *L*^2^-norm, and node persistence, with the phenotypic scores available for each cognitive or behavioral test. We employed Spearman correlation as certain test scores are ordinal rather than continuous. Finally, we adjusted the p-values associated with the estimated correlations using FDR correction. The FDR correction was implemented separately for p-values of each cognitive or behavioral test. For example, if a given cognitive or behavioral test contains more than one phenotypic score, we performed the FDR correction across all associated scores.

For the ABIDE-I dataset, the analysis was conducted solely for the ASD group. We opted for two clinical scores according to the Autism Diagnostic Interview-Revised (ADI-R) scoring criteria [116], namely ADI-R verbal and ADI-R social. Among all the potential clinical scores, the ADI-R scores are particularly noteworthy for two main reasons. First, they are available for the largest segment of the ASD group (275 participants). Second, the ADI-R social and verbal scores provide an effective measure of symptom severity in autism [117]. We calculated the Spearman correlation between PH-based measures and clinical scores. However, no FDR correction was applied at the global and mesoscopic scales since only two scores were considered.

### Estimating overlap between brain regions identified by ROI-level analysis and non-invasive stimulation studies

We conducted a systematic comparison between the ROIs showing significant between-group differences as identified by local topological measures and those identified from non-invasive brain stimulation (NIBS) studies. We focused on three NIBS modalities, namely transcranial direct current stimulation (tDCS), transcranial alternating current stimulation (tACS), and transcranial magnetic stimulation (TMS).

To compare with brain regions exhibiting age-related changes in functional connectivity, we focused on target regions whose non-invasive stimulation led to enhanced motor performance in healthy elderly individuals. To compare with brain regions exhibiting ASD-related changes in functional connectivity, we focused on target regions whose non-invasive stimulation led to improvement of symptoms associated with ASD. We evaluated the overlap of these target regions with those identified by node persistence, node frequency, and nodal PSS. These overlaps allowed us to assess whether local topological measures of resting-state functional connectivity could identify brain regions whose non-invasive stimulation yields functional benefits in healthy aging and ASD.

The target regions of NIBS experiments were acquired based on a systematic literature review carried out in earlier studies by some of us [67, 69]. These target regions, which originally correspond to regions in the Brodmann atlas [118], were mapped to regions in the Schaefer atlas. This mapping is available at https://github.com/asamallab/Curvature-FCN-ASD.

## ACKNOWLEDGMENTS

The authors thank Kelin Xia for insightful discussions. A.S. acknowledges funding from the Department of Atomic Energy, Government of India (via the Apex project to The Institute of Mathematical Sciences (IMSc), Chennai) and funding from the Max Planck Society, Germany (through the award of a Max Planck Partner Group).

## AUTHOR CONTRIBUTION

Designed the research: M.M., Y.Y., J.J., A.S.; Performed the research: M.M., Y.Y., J.J., A.S.; Performed the computations: M.M.; Wrote the paper: M.M., Y.Y., J.J., A.S.

## DATA AND CODE AVAILABILITY

The functional connectivity matrices considered in our study, as well as all the code used for the analysis and to reproduce the results in this manuscript, are publicly accessible through the GitHub repository: https://github.com/asamallab/NodePersistence_PH_FCM.

## References

[1] K.J. Friston, C.D. Frith, P.F. Liddle, and R.S. Frackowiak. Functional connectivity: the principal-component analysis of large (PET) data sets. Journal of Cerebral Blood Flow & Metabolism, 13(1):5–14, 1993.

[2] S. Ogawa, T.M. Lee, A.R. Kay, and D.W. Tank. Brain magnetic resonance imaging with contrast dependent on blood oxygenation. Proceedings of the National Academy of Sciences, 87(24):9868–9872, 1990.

[3] N.K. Logothetis. What we can do and what we cannot do with fMRI. Nature, 453(7197):869–878, 2008.

[4] J. Hlinka, M. Paluš, M. Vejmelka, D. Mantini, and M. Corbetta. Functional connectivity in resting-state fMRI: is linear correlation sufficient? NeuroImage, 54(3):2218–2225, 2011.

[5] B. Biswal, F. Z. Yetkin, V.M. Haughton, and J.S. Hyde. Functional connectivity in the motor cortex of resting human brain using echo-planar MRI. Magnetic Resonance in Medicine, 34(4):537–541, 1995.

[6] M.E. Raichle, A.M. MacLeod, A.Z. Snyder, W.J. Powers, D.A. Gusnard, and G.L. Shulman. A default mode of brain function. Proceedings of the National Academy of Sciences, 98(2):676–682, 2001.

[7] M.D. Greicius, B. Krasnow, A.L. Reiss, and V. Menon. Functional connectivity in the resting brain: A network analysis of the default mode hypothesis. Proceedings of the National Academy of Sciences, 100(1):253–258, 2003.

[8] S.M Smith, P.T. Fox, K.L. Miller, D.C. Glahn, P.M. Fox, C.E. Mackay, N. Filippini, K.E. Watkins, R. Toro, A.R. Laird, et al. Correspondence of the brain’s functional architecture during activation and rest. Proceedings of the National Academy of Sciences, 106(31):13040–13045, 2009.

[9] M.D. Fox and M.E. Raichle. Spontaneous fluctuations in brain activity observed with functional magnetic resonance imaging. Nature Reviews Neuroscience, 8(9):700–711, 2007.

[10] B.T.T. Yeo, F.M. Krienen, J. Sepulcre, M.R. Sabuncu, D. Lashkari, M. Hollinshead, J.L. Roffman, J.W. Smoller, L. Zöllei, J.R. Polimeni, et al. The organization of the human cerebral cortex estimated by intrinsic functional connectivity. Journal of Neurophysiology, 106(3), 2011.

[11] C. Grady. The cognitive neuroscience of ageing. Nature Reviews Neuroscience, 13(7):491–505, 2012.

[12] D. Tromp, A. Dufour, S. Lithfous, T. Pebayle, and O. Després. Episodic memory in normal aging and Alzheimer disease: Insights from imaging and behavioral studies. Ageing research reviews, 24:232–262, 2015.

[13] L. Nyberg, A. Salami, M. Andersson, J. Eriksson, G. Kalpouzos, K. Kauppi, J. Lind, S. Pudas, J. Persson, and L.-G. Nilsson. Longitudinal evidence for diminished frontal cortex function in aging. Proceedings of the National Academy of Sciences, 107(52):22682–22686, 2010.

[14] C. Lord, T.S. Brugha, T. Charman, J. Cusack, G. Dumas, T. Frazier, E.J.H. Jones, R.M. Jones, A. Pickles, M.W. State, et al. Autism spectrum disorder. Nature Reviews Disease Primers, 6(1):5, 2020.

[15] National Institute of Neurological Disorders and Stroke. Autism Spectrum Disorder Fact Sheet, 2020.

[16] S. Kristen, F. Rossmann, and B. Sodian. Theory of own mind and autobiographical memory in adults with ASD. Research in Autism Spectrum Disorders, 8(7):827–837, 2014.

[17] A. Habib, L. Harris, F. Pollick, and C. Melville. A meta-analysis of working memory in individuals with autism spectrum disorders. PloS ONE, 14(4):e0216198, 2019.

[18] C.J. Zampella, L.A.L. Wang, M. Haley, A.G. Hutchinson, and A. de Marchena. Motor Skill Differences in Autism Spectrum Disorder: a Clinically Focused Review. Current Psychiatry Reports, 23(10):64, 2021.

[19] N.K. Arora, M.K.C. Nair, S. Gulati, V. Deshmukh, A. Mohapatra, D. Mishra, V. Patel, R.M. Pandey, B.C. Das, G. Divan, et al. Neurodevelopmental disorders in children aged 2–9 years: Population-based burden estimates across five regions in India. PLoS Medicine, 15(7):e1002615, 2018.

[20] M.J. Maenner, K.A. Shaw, J. Baio, A. Washington, M. Patrick, M. DiRienzo, D.L. Christensen, L.D. Wiggins, S. Pettygrove, J.G. Andrews, et al. Prevalence of autism spectrum disorder among children aged 8 years – autism and developmental disabilities monitoring network, 11 sites, United States, 2016. MMWR. Surveillance Summaries, 69(4):1–12, 2020.

[21] D. Fein, M. Barton, I.-M. Eigsti, E. Kelley, L. Naigles, R.T. Schultz, M. Stevens, M. Helt, A. Orinstein, M. Rosenthal, et al. Optimal outcome in individuals with a history of autism. Journal of Child Psychology and Psychiatry, 54(2):195–205, 2013.

[22] R.N. Spreng, M. Wojtowicz, and C.L. Grady. Reliable differences in brain activity between young and old adults: A quantitative meta-analysis across multiple cognitive domains. Neuroscience & Biobehavioral Reviews, 34(8):1178–1194, 2010.

[23] N.D. Woodward and C.J. Cascio. Resting-State Functional Connectivity in Psychiatric Disorders. JAMA Psychiatry, 72(8):743–744, 2015.

[24] S. Solso, R. Xu, J. Proudfoot, D.J. Hagler, K. Campbell, V. Venkatraman, C.C. Barnes, C. Ahrens-Barbeau, K. Pierce, A. Dale, et al. Diffusion Tensor Imaging Provides Evidence of Possible Axonal Overconnectivity in Frontal Lobes in Autism Spectrum Disorder Toddlers. Biological Psychiatry, 79(8):676–684, 2016.

[25] F. Chazal and B. Michel. An introduction to topological data analysis: fundamental and practical aspects for data scientists. Frontiers in Artificial Intelligence, 4:667963, 2021.

[26] H. Edelsbrunner, D. Letscher, and A. Zomorodian. Topological Persistence and Simplification. Discrete & Computational Geometry, 28(4):511–533, 2002.

[27] A. Patania, F. Vaccarino, and G. Petri. Topological analysis of data. EPJ Data Science, 6(1):7, 2017.

[28] J.R. Munkres. Elements of Algebraic Topology. CRC press, 2018.

[29] R. Ghrist. Barcodes: The persistent topology of data. Bulletin of the American Mathematical Society, 45(1):61–75, 2008.

[30] H. Edelsbrunner and J. Harer. Persistent Homology – a Survey. Contemporary Mathematics, 453(26):257–282, 2008.

[31] G. Carlsson. Topology and data. Bulletin of the American Mathematical Society, 46(2):255–308, 2009.

[32] H. Edelsbrunner and J. Harer. Computational Topology: An Introduction. American Mathematical Society, 2010.

[33] J.M. Chan, G. Carlsson, and R. Rabadan. Topology of viral evolution. Proceedings of the National Academy of Sciences, 110(46):18566–18571, 2013.

[34] C.M. Topaz, L. Ziegelmeier, and T. Halverson. Topological data analysis of biological aggregation models. PLoS ONE, 10(5):e0126383, 2015.

[35] P.G. Cámara, A.J. Levine, and R. Rabadán. Inference of Ancestral Recombination Graphs through Topological Data Analysis. PLoS Computational Biology, 12(8):e1005071, 2016.

[36] K. Xia, Z. Li, and L. Mu. Multiscale Persistent Functions for Biomolecular Structure Characterization. Bulletin of Mathematical Biology, 80:1–31, 2018.

[37] M. Gidea. Topological Data Analysis of Critical Transitions in Financial Networks. In 3rd International Winter School and Conference on Network Science: NetSci-X 2017, pages 47–59, 2017.

[38] M. Gidea and Y. Katz. Topological data analysis of financial time series: Landscapes of crashes. Physica A: Statistical Mechanics and its Applications, 491:820–834, 2018.

[39] H. Guo, X. Zhao, H. Yu, and X. Zhang. Analysis of global stock markets’ connections with emphasis on the impact of COVID-19. Physica A: Statistical Mechanics and its Applications, 569:125774, 2021.

[40] S. Kulkarni, H.K. Pharasi, S. Vijayaraghavan, S. Kumar, A. Chakraborti, and A. Samal. Investigation of Indian stock markets using topological data analysis and geometry-inspired network measures. Physica A: Statistical Mechanics and its Applications, 643:129785, 2024.

[41] M. Kramar, A. Goullet, L. Kondic, and K. Mischaikow. Persistence of force networks in compressed granular media. Physical Review E, 87(4):042207, 2013.

[42] I. Donato, M. Gori, M. Pettini, G. Petri, S.D. Nigris, R. Franzosi, and F. Vaccarino. Persistent homology analysis of phase transitions. Physical Review E, 93(5):052138, 2016.

[43] S. Heydenreich, B. Brück, and J. Harnois-Déraps. Persistent homology in cosmic shear: Constraining parameters with topological data analysis. Astronomy & Astrophysics, 648:A74, 2021.

[44] H. Adams, T. Emerson, M. Kirby, R. Neville, C. Peterson, P. Shipman, S. Chepushtanova, E. Hanson, F. Motta, and L. Ziegelmeier. Persistence images: a stable vector representation of persistent homology. The Journal of Machine Learning Research, 18(1):218–252, 2017.

[45] J. Townsend, C.P. Micucci, J.H. Hymel, V. Maroulas, and K.D. Vogiatzis. Representation of molecular structures with persistent homology for machine learning applications in chemistry. Nature Communications, 11(1):3230, 2020.

[46] C.S. Pun, S.X. Lee, and K. Xia. Persistent-homology-based machine learning: a survey and a comparative study. Artificial Intelligence Review, 55(7):5169–5213, 2022.

[47] R. Vandaele, G.A. Nervo, and O. Gevaert. Topological image modification for object detection and topological image processing of skin lesions. Scientific Reports, 10(1):21061, 2020.

[48] H. Edelsbrunner. Persistent Homology in Image Processing. In Graph-Based Representations in Pattern Recognition, pages 182–183, 2013.

[49] G. Petri, P. Expert, F. Turkheimer, R. Carhart-Harris, D. Nutt, P.J. Hellyer, and F. Vaccarino. Homological scaffolds of brain functional networks. Journal of The Royal Society Interface, 11(101):20140873, 2014.

[50] L.-D. Lord, P. Expert, H.M. Fernandes, G. Petri, T.J. Van Hartevelt, F. Vaccarino, G. Deco, F. Turkheimer, and M.L. Kringelbach. Insights into Brain Architectures from the Homological Scaffolds of Functional Connectivity Networks. Frontiers in Systems Neuroscience, 10:85, 2016.

[51] C. Giusti, R. Ghrist, and D.S. Bassett. Two’s company, three (or more) is a simplex. Journal of Computational Neuroscience, 41:1–14, 2016.

[52] P. Bendich, J.S. Marron, E. Miller, A. Pieloch, and S. Skwerer. Persistent homology analysis of brain artery trees. The Annals of Applied Statistics, 10(1):198–218, 2016.

[53] M.K. Chung, H. Lee, A. DiChristofano, H. Ombao, and V. Solo. Exact topological inference of the resting-state brain networks in twins. Network Neuroscience, 3(3):674–694, 2019.

[54] L. Kuang, X. Han, K. Chen, R.J. Caselli, E.M. Reiman, Y. Wang, and Alzheimer’s Disease Neuroimaging Initiative. A concise and persistent feature to study brain resting-state network dynamics: Findings from the Alzheimer’s Disease Neuroimaging Initiative. Human Brain Mapping, 40(4):1062–1081, 2019.

[55] D. Liang, S. Xia, X. Zhang, and W. Zhang. Analysis of Brain Functional Connectivity Neural Circuits in Children With Autism Based on Persistent Homology. Frontiers in Human Neuroscience, 15:745671, 2021.

[56] A.T. Jafadideh and B.M. Asl. Topological analysis of brain dynamics in autism based on graph and persistent homology. Computers in Biology and Medicine, 150:106202, 2022.

[57] J. Xing, J. Jia, X. Wu, and L. Kuang. A Spatiotemporal Brain Network Analysis of Alzheimer’s Disease Based on Persistent Homology. Frontiers in Aging Neuroscience, 14:788571, 2022.

[58] W. Zhang, S. Xia, X. Tang, X. Zhang, D. Liang, and Y. Wang. Topological analysis of functional connectivity in Parkinson’s disease. Frontiers in Neuroscience, 17:1236128, 2023.

[59] H. Ryu, C. Habeck, Y. Stern, and S. Lee. Persistent homology-based functional connectivity and its association with cognitive ability: Life-span study. Human Brain Mapping, 44(9):3669–3683, 2023.

[60] M. Mijangos, L. Pacheco, A. Bravetti, N. González-García, P. Padilla, and R. Velasco-Segura. Persistent homology reveals robustness loss in inhaled substance abuse rs-fMRI networks. PloS ONE, 19(9):e0310165, 2024.

[61] M.K. Chung, S.-G. Huang, I.C. Carroll, V.D. Calhoun, and H.H. Goldsmith. Topological state-space estimation of functional human brain networks. PLOS Computational Biology, 20(5):e1011869, 2024.

[62] M. Aggarwal and V. Periwal. Tight basis cycle representatives for persistent homology of large biological data sets. PLOS Computational Biology, 19(5):e1010341, 2023.

[63] P. Bendich, E. Gasparovic, J. Harer, R. Izmailov, and L. Ness. Multi-scale local shape analysis and feature selection in machine learning applications. In 2015 International Joint Conference on Neural Networks (IJCNN), pages 1–8, 2015.

[64] B.T. Fasy and B. Wang. Exploring persistent local homology in topological data analysis. In 2016 IEEE International Conference on Acoustics, Speech and Signal Processing (ICASSP), pages 6430–6434, 2016.

[65] N. Nguyen, T. Hou, E. Amico, J. Zheng, H. Huang, A.D. Kaplan, G. Petri, J. Goñi, R. Kaufmann, Y. Zhao, et al. Volume-Optimal Persistence Homological Scaffolds of Hemodynamic Networks Covary with MEG Theta-Alpha Aperiodic Dynamics. In Medical Image Computing and Computer Assisted Intervention – MICCAI 2024, pages 519–529, 2024.

[66] A. Babayan, M. Erbey, D. Kumral, J.D. Reinelt, A.M.F. Reiter, J. Röbbig, H.L. Schaare, M. Uhlig, A. Anwander, P.-L. Bazin, et al. A mind-brain-body dataset of MRI, EEG, cognition, emotion, and peripheral physiology in young and old adults. Scientific Data, 6(1):180308, 2019.

[67] Y. Yadav, P. Elumalai, N. Williams, J. Jost, and A. Samal. Discrete Ricci curvatures capture age-related changes in human brain functional connectivity networks. Frontiers in Aging Neuroscience, 15:1120846, 2023.

[68] A. Di Martino, C.G. Yan, Q. Li, E. Denio, F.X. Castellanos, K. Alaerts, J.S. Anderson, M. Assaf, S.Y. Bookheimer, M. Dapretto, et al. The autism brain imaging data exchange: towards a large-scale evaluation of the intrinsic brain architecture in autism. Molecular Psychiatry, 19(6):659–667, 2014.

[69] P. Elumalai, Y. Yadav, N. Williams, E. Saucan, J. Jost, and A. Samal. Graph Ricci curvatures reveal atypical functional connectivity in autism spectrum disorder. Scientific Reports, 12(1):8295, 2022.

[70] T. Yarkoni, R.A. Poldrack, T.E. Nichols, D.C. Van Essen, and T.D. Wager. Large-scale automated synthesis of human functional neuroimaging data. Nature Methods, 8(8):665–670, 2011.

[71] N. Williams, S.H. Wang, G. Arnulfo, L. Nobili, S. Palva, and J.M. Palva. Modules in connectomes of phase-synchronization comprise anatomically contiguous, functionally related regions. NeuroImage, 272:120036, 2023.

[72] A. Schaefer, R. Kong, E.M. Gordon, T.O. Laumann, X.-N. Zuo, A.J. Holmes, S.B. Eickhoff, and B.T.T. Yeo. Local-Global Parcellation of the Human Cerebral Cortex from Intrinsic Functional Connectivity MRI. Cerebral Cortex, 28(9):3095–3114, 2018.

[73] R.N. Mantegna. Hierarchical structure in financial markets. The European Physical Journal B - Condensed Matter and Complex Systems, 11(1):193–197, 1999.

[74] J. Qian, I. Diez, L. Ortiz-Terán, C. Bonadio, T. Liddell, J. Goñi, and J. Sepulcre. Positive Connectivity Predicts the Dynamic Intrinsic Topology of the Human Brain Network. Frontiers in Systems Neuroscience, 12:38, 2018.

[75] H. Chintakunta, T. Gentimis, R. Gonzalez-Diaz, M.-J. Jimenez, and H. Krim. An entropy-based persistence barcode. Pattern Recognition, 48(2):391–401, 2015.

[76] P. Bubenik. The Persistence Landscape and Some of Its Properties. In Topological Data Analysis, pages 97–117, 2020.

[77] K.K. Yuen. The two-sample trimmed t for unequal population variances. Biometrika, 61(1):165–170, 1974.

[78] M. Xia, J. Wang, and Y. He. BrainNet Viewer: A Network Visualization Tool for Human Brain Connectomics. PloS ONE, 8(7):e68910, 2013.

[79] H. Adams, A. Tausz, and M. Vejdemo-Johansson. JavaPlex: A research software package for persistent (co)homology. In Mathematical Software–ICMS 2014, pages 129–136. Springer Berlin Heidelberg, 2014.

[80] M. Krassowski. ComplexUpset, 2020.

[81] R.D. Seidler, J.A. Bernard, T.B. Burutolu, B.W. Fling, M.T. Gordon, J.T. Gwin, Y. Kwak, and D.B. Lipps. Motor control and aging: Links to age-related brain structural, functional, and biochemical effects. Neuroscience & Biobehavioral Reviews, 34(5):721–733, 2010.

[82] J. Cirillo. Physical activity, motor performance and skill learning: a focus on primary motor cortex in healthy aging. Experimental Brain Research, 239(12):3431–3438, 2021.

[83] R.D. Seidler, J.L. Alberts, and G.E. Stelmach. Changes in Multi-Joint Performance with Age. Motor Control, 6(1):19–31, 2002.

[84] J.L. Contreras-Vidal, H.-L. Teulings, and G.E. Stelmach. Elderly subjects are impaired in spatial coordination in fine motor control. Acta Psychologica, 100(1):25–35, 1998.

[85] V.D. Buckles. Age-Related Slowing, pages 73–87. Springer Netherlands, 1993.

[86] M. Schmitter-Edgecombe, M. Vesneski, and D.W.R. Jones. Aging and Word-Finding: A Comparison of Spontaneous and Constrained Naming Tests. Archives of Clinical Neuropsychology, 15(6):479–493, 2000.

[87] M. Mather. The emotion paradox in the aging brain. Annals of the New York Academy of Sciences, 1251(1):33–49, 2012.

[88] R.R. Edwards and R.B. Fillingim. Age-Associated Differences in Responses to Noxious Stimuli. The Journals of Gerontology: Series A, 56(3):M180–M185, 2001.

[89] X. Ming, M. Brimacombe, and G.C. Wagner. Prevalence of motor impairment in autism spectrum disorders. Brain and Development, 29(9):565–570, 2007.

[90] A. Pickles, E. Simonoff, G. Conti-Ramsden, M. Falcaro, Z. Simkin, T. Charman, S. Chandler, T. Loucas, and G. Baird. Loss of language in early development of autism and specific language impairment. Journal of Child Psychology and Psychiatry, 50(7):843–852, 2009.

[91] S.E. Weismer, C. Lord, and A. Esler. Early language patterns of toddlers on the autism spectrum compared to toddlers with developmental delay. Journal of Autism and Developmental Disorders, 40(10):1259–1273, 2010.

[92] C. Kasari and S. Patterson. Interventions Addressing Social Impairment in Autism. Current Psychiatry Reports, 14(6):713–725, 2012.

[93] A. Senju. Spontaneous Theory of Mind and Its Absence in Autism Spectrum Disorders. The Neuroscientist, 18(2):108–113, 2012.

[94] M. Solomon, J.B. McCauley, A.-M. Iosif, C.S. Carter, and J.D. Ragland. Cognitive control and episodic memory in adolescents with autism spectrum disorders. Neuropsychologia, 89:31–41, 2016.

[95] A.N. Bhat. Motor impairment increases in children with autism spectrum disorder as a function of social communication, cognitive and functional impairment, repetitive behavior severity, and comorbid diagnoses: A SPARK study report. Autism Research, 14(1):202–219, 2021.

[96] L. Vietoris. Über den höheren Zusammenhang kompakter Räume und eine Klasse von zusammenhangstreuen Abbildungen. Mathematische Annalen, 97(1):454–472, 1927.

[97] S. Whitfield-Gabrieli and A. Nieto-Castanon. Conn: A Functional Connectivity Toolbox for Correlated and Anticorrelated Brain Networks. Brain Connectivity, 2(3):125–141, 2012.

[98] A. Hatcher. Algebraic Topology. Cambridge University Press, 2002.

[99] N. Otter, M.A. Porter, U. Tillmann, P. Grindrod, and H.A. Harrington. A roadmap for the computation of persistent homology. EPJ Data Science, 6(1):17, 2017.

[100] S. Dantchev and I. Ivrissimtzis. Efficient construction of the Čech complex. Computers & Graphics, 36(6):708–713, 2012.

[101] M. Kerber and R. Sharathkumar. Approximate Čech Complex in Low and High Dimensions. In Algorithms and Computation, pages 666–676. Springer Berlin Heidelberg, 2013.

[102] G. Chen, G. Chen, C. Xie, and S.-J. Li. Negative Functional Connectivity and Its Dependence on the Shortest Path Length of Positive Network in the Resting-State Human Brain. Brain Connectivity, 1(3):195–206, 2011.

[103] V. Robins. Towards computing homology from finite approximations. In Topology Proceedings, volume 24, pages 503–532, 1999.

[104] A. Zomorodian and G. Carlsson. Computing Persistent Homology. Discrete & Computational Geometry, 33(2):249–274, 2005.

[105] M. Rucco, F. Castiglione, E. Merelli, and M. Pettini. Characterisation of the Idiotypic Immune Network Through Persistent Entropy. In Proceedings of ECCS 2014, pages 117–128, 2016.

[106] N. Atienza, R. Gonzalez-Díaz, and M. Soriano-Trigueros. On the stability of persistent entropy and new summary functions for topological data analysis. Pattern Recognition, 107:107509, 2020.

[107] D. Cohen-Steiner, H. Edelsbrunner, and J. Harer. Stability of persistence diagrams. In Proceedings of the Twenty-First Annual Symposium on Computational Geometry, pages 263–271, 2005.

[108] Y. Mileyko, S. Mukherjee, and J. Harer. Probability measures on the space of persistence diagrams. Inverse Problems, 27(12):124007, 2011.

[109] K. Turner, Y. Mileyko, S. Mukherjee, and J. Harer. Fréchet Means for Distributions of Persistence Diagrams. Discrete & Computational Geometry, 52(1):44–70, 2014.

[110] P. Bubenik. Statistical topological data analysis using persistence landscapes. Journal of Machine Learning Research, 16(1):77–102, 2015.

[111] C. Maria, J.-D. Boissonnat, M. Glisse, and M. Yvinec. The Gudhi Library: Simplicial Complexes and Persistent Homology. In Mathematical Software–ICMS 2014, pages 167–174, 2014.

[112] M. Guerra, A.D. Gregorio, U. Fugacci, G. Petri, and F. Vaccarino. Homological scaffold via minimal homology bases. Scientific Reports, 11(1):5355, 2021.

[113] Y. Benjamini and Y. Hochberg. Controlling the false discovery rate: a practical and powerful approach to multiple testing. Journal of the Royal Statistical Society: Series B (Methodological), 57(1):289–300, 1995.

[114] P. Virtanen, R. Gommers, T.E. Oliphant, M. Haberland, T. Reddy, D. Cournapeau, E. Burovski, P. Peterson, W. Weckesser, J. Bright, et al. SciPy 1.0: fundamental algorithms for scientific computing in Python. Nature Methods, 17(3):261–272, 2020.

[115] S. Seabold and J. Perktold. Statsmodels: Econometric and Statistical Modeling with Python. In Proceedings of the 9th Python in Science Conference, 2010.

[116] C. Lord, M. Rutter, and A. Le Couteur. Autism Diagnostic Interview-Revised: A revised version of a diagnostic interview for caregivers of individuals with possible pervasive developmental disorders. Journal of Autism and Developmental Disorders, 24(5):659–685, 1994.

[117] J. Lefort-Besnard, K. Vogeley, L. Schilbach, G. Varoquaux, B. Thirion, G. Dumas, and D. Bzdok. Patterns of autism symptoms: hidden structure in the ADOS and ADI-R instruments. Translational Psychiatry, 10(1):257, 2020.

[118] M. Strotzer. One century of brain mapping using Brodmann areas. Clinical Neuroradiology, 19(3):179–186, 2009.

