## Supplementary Text for "Node persistence from topological data analysis reveals changes in brain functional connectivity"

### SUPPLEMENTARY TEXT
### for
### Node persistence from topological data analysis reveals changes in brain functional connectivity

Madhumita Mondal,[1,2] Yasharth Yadav,[3] Jürgen Jost,[4,5,6,7,*] and Areejit Samal[1,2,*]

[1]*The Institute of Mathematical Sciences (IMSc), Chennai 600113, India*
[2]*Homi Bhabha National Institute (HBNI), Mumbai 400094, India*
[3]*School of Physical and Mathematical Sciences, Nanyang Technological University, 637371, Singapore*
[4]*Max Planck Institute for Mathematics in the Sciences, 04103 Leipzig, Germany*
[5]*Max Planck Institute for Human Cognitive and Brain Sciences, 04103 Leipzig, Germany*
[6]*Center for Scalable Data Analytics and Artificial Intelligence, Dresden/Leipzig, 04107 Leipzig, Germany*
[7]*Santa Fe Institute for the Sciences of Complexity, Santa Fe, NM 87501, USA*

**Comparison with existing local measures based on persistent homology**

The concept of a homological scaffold was introduced by Petri *et al.* as a secondary network summarizing one-dimensional holes identified through PH [1]. Nodal Persistence Scaffold Strength (PSS) was introduced to evaluate the centrality of a node to measure the significance of a region that contributes to these one-dimensional holes or cycles [2]. In this study, we compared nodal PSS against our newly proposed local metrics: node persistence and node frequency, since all of these metrics characterize the structure of one-dimensional holes. However, generating homological scaffolds is computationally expensive for larger point sets, and particularly not scalable for simplicial complexes with 200 points. Therefore, for a fair comparison between nodal PSS and our proposed metrics, we investigated the region of interest (ROI)-level changes in functional connectivity using functional connectivity (FC) matrices at the resting-state networks (RSNs)-level. This approach targets intra-RSN topological patterns through FC matrices that are specific to a given RSN.

*RSN-to-local level analysis*

We computed node persistence and node frequency using Rips complexes constructed from submatrices of the FC Matrix at the level of individual RSNs. A two-tailed two-sample t-test was utilized to detect statistical differences between the groups, and FDR correction was applied independently for each RSN.

In the MPI-LEMON dataset, 59 ROIs exhibit significant differences ($p < 0.05$, FDR-corrected) between the young and elderly groups considering node persistence. These ROIs are distributed among six RSNs: visual (1), somatomotor (24), dorsal attention (5), salience/ventral attention (1), control (1), and default (27) networks. All of these ROIs show higher node persistence values in young individuals compared to elderly individuals. In the ABIDE-I dataset, 53 ROIs exhibit significant differences ($p < 0.05$, FDR-corrected) between the ASD and TD groups via node persistence. These ROIs are distributed among the three RSNs: somatomotor (10), salience/ventral attention (5), and default (38) networks. All the ROIs show higher node persistence values for individuals with ASD compared to TD individuals.

Node frequency reveals 40 and 35 ROIs with significant between-group differences ($p < 0.05$, FDR-corrected) for the MPI-LEMON and ABIDE-I datasets, respectively. In the MPI-LEMON dataset, 40 ROIs are distributed in the RSNs as follows: somatomotor (15), dorsal attention (4), salience/ventral attention (3), control (2), and default (16) networks. Moreover, 37 out of these 40 ROIs identified by node frequency are also identified via node persistence. In the ABIDE-I dataset, 35 ROIs are distributed in three RSNs as follows: somatomotor (8), salience/ventral attention (8), and default (19) networks. However, 31 out of these 35 ROIs overlap with those found via node persistence. All the ROIs show higher node frequency values in young individuals than in elderly individuals within the MPI-LEMON dataset, and in ASD individuals compared to TD individuals within the ABIDE-I dataset.

In the MPI-LEMON dataset, nodal PSS detects 58 ROIs with significant between-group differences ($p < 0.05$, FDR-corrected) between young and elderly individuals. These include ROIs from six RSNs: visual (2), somatomotor (26), dorsal attention (5), salience/ventral attention (7), control (2), and default (16) networks. Of these 58 ROIs,

47 were also identified via node persistence, and 36 were identified via node frequency. However, in this case, 37 out of 58 ROIs show higher nodal PSS values in young individuals than in elderly individuals. In the ABIDE-I dataset, nodal PSS detects 45 ROIs from the three RSNs, somatomotor (7), salience/ventral attention (14), and default (24), with significant differences between the ASD and TD group ($p < 0.05$, FDR-corrected). Of these 45 ROIs, 35 are also identified through node persistence, and 31 ROIs are identified through node frequency. In this case, most of the ROIs (42 out of 45) display higher nodal PSS values in the ASD group compared to the TD group.

All ROI-level differences identified using node persistence, node frequency, and nodal PSS for both datasets are listed in Supplementary Table S10. Additionally, Supplementary Table S11 provides the group-wise averages of node persistence, node frequency, and nodal PSS for each of the 200 ROIs across both datasets, along with the corresponding FDR-corrected $p$-values.

##### *Linking RSN-to-local level analysis with non-invasive brain stimulation outcomes*

We investigated the relevance of ROI-level differences in PH-based local measures with the existing literature on non-invasive brain stimulation (NIBS) in healthy elderly individuals and individuals with ASD. As mentioned in the Results section in the main text, target regions that have been shown to enhance motor performance in healthy elderly individuals [3] correspond to 42 ROIs in the Schaefer atlas. Furthermore, 31 Schaefer ROIs show evidence for improving behavioral or cognitive symptoms associated with ASD [4].

The UpSet plots in Supplementary Fig. S6 illustrate the number of ROIs associated with clinical improvement and their overlaps with ROIs detected by node persistence, node frequency, and nodal PSS, for both the MPI-LEMON and ABIDE-I datasets. For the MPI-LEMON dataset, out of the 42 clinically relevant ROIs identified by NIBS, 19 are captured by at least one of the topology-based local measures, and 10 are identified by all three measures. In particular, node persistence reveals 15 clinically relevant ROIs spanning three RSNs, node frequency identifies 13 ROIs spanning five RSNs, and nodal PSS reveals 18 ROIs spanning five RSNs (see Supplementary Fig. S6(a)). For the ABIDE-I dataset, out of the 31 clinically relevant ROIs identified by NIBS, 11 are captured by at least one of the three topology-based local measures, and those are distributed in three RSNs: somatomotor, salience/ventral attention, and default networks. Among these 11 ROIs, 4 ROIs are uniquely identified by node persistence, whereas 2 are uniquely identified by nodal PSS. Specifically, node persistence detects 9 clinically relevant ROIs, node frequency identifies 3, and nodal PSS identifies 7 ROIs (see Supplementary Fig. S6(b) and Supplementary Table S9).

---

[1] G. Petri, P. Expert, F. Turkheimer, R. Carhart-Harris, D. Nutt, P.J. Hellyer, and F. Vaccarino. Homological scaffolds of brain functional networks. *Journal of The Royal Society Interface*, 11(101):20140873, 2014.

[2] L.-D. Lord, P. Expert, H.M. Fernandes, G. Petri, T.J. Van Hartevelt, F. Vaccarino, G. Deco, F. Turkheimer, and M.L. Kringelbach. Insights into Brain Architectures from the Homological Scaffolds of Functional Connectivity Networks. *Frontiers in Systems Neuroscience*, 10:85, 2016.

[3] Y. Yadav, P. Elumalai, N. Williams, J. Jost, and A. Samal. Discrete Ricci curvatures capture age-related changes in human brain functional connectivity networks. *Frontiers in Aging Neuroscience*, 15:1120846, 2023.

[4] P. Elumalai, Y. Yadav, N. Williams, E. Saucan, J. Jost, and A. Samal. Graph Ricci curvatures reveal atypical functional connectivity in autism spectrum disorder. *Scientific Reports*, 12(1):8295, 2022.
