## Supplementary Figure for "Node persistence from topological data analysis reveals changes in brain functional connectivity"

### SUPPLEMENTARY FIGURES
### for
### Node persistence from topological data analysis reveals changes in brain functional connectivity


Madhumita Mondal,[1,2] Yasharth Yadav,[3] Jürgen Jost,[4,5,6,7,*] and Areejit Samal[1,2,*]

[1] *The Institute of Mathematical Sciences (IMSc), Chennai 600113, India*
[2] *Homi Bhabha National Institute (HBNI), Mumbai 400094, India*
[3] *School of Physical and Mathematical Sciences, Nanyang Technological University, 637371, Singapore*
[4] *Max Planck Institute for Mathematics in the Sciences, 04103 Leipzig, Germany*
[5] *Max Planck Institute for Human Cognitive and Brain Sciences, 04103 Leipzig, Germany*
[6] *Center for Scalable Data Analytics and Artificial Intelligence, Dresden/Leipzig, 04107 Leipzig, Germany*
[7] *Santa Fe Institute for the Sciences of Complexity, Santa Fe, NM 87501, USA*

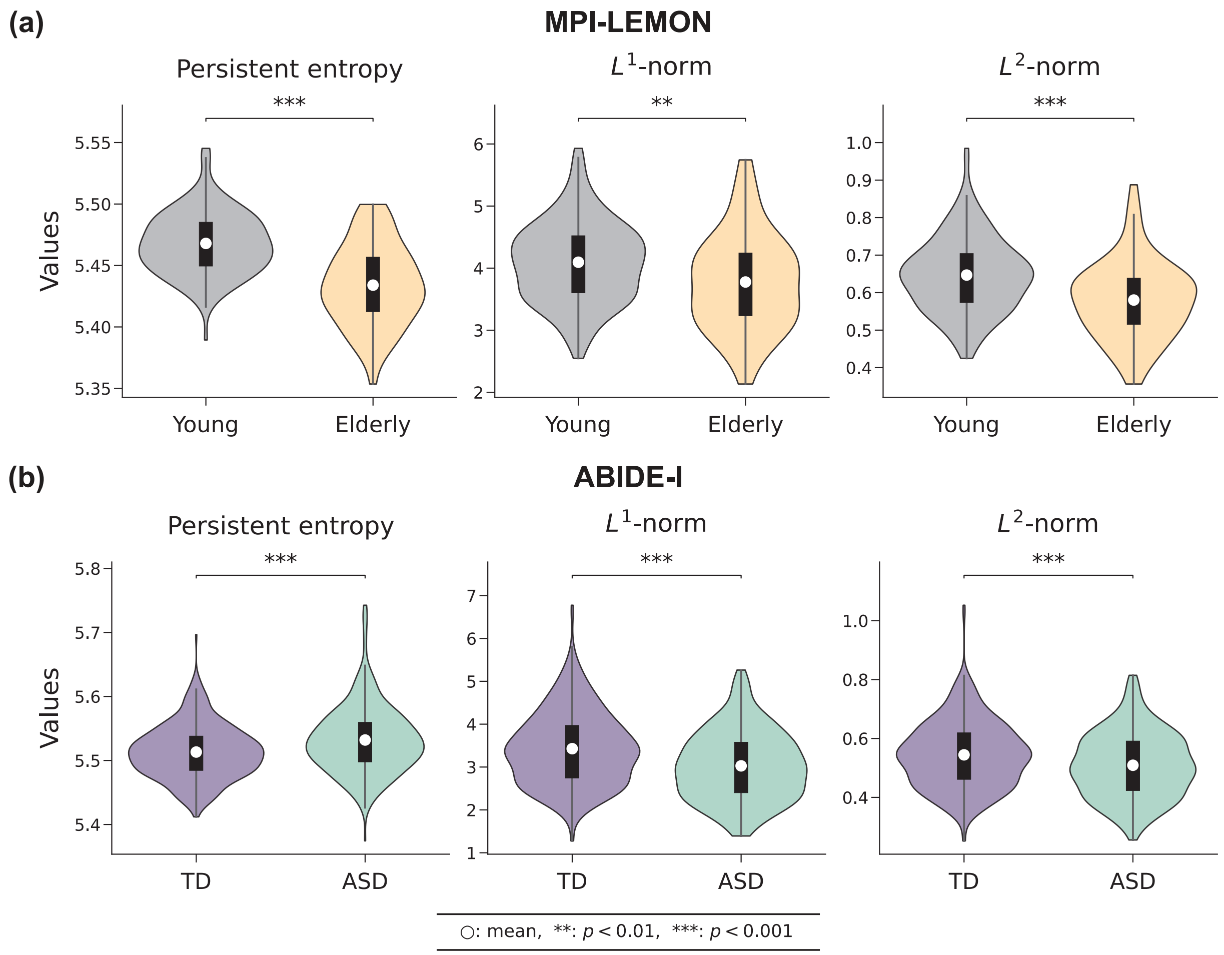


FIG. S1. **Brain-wide differences between the groups as identified by three global measures: persistent entropy of the persistence barcodes, $L^1$-norm, and $L^2$-norm of the persistent landscape, considering all the correlations from functional connectivity matrices.** (a) MPI-LEMON dataset: violin plots corresponding to 153 young and 72 elderly individuals across three global measures. The mean values of all three measures are significantly higher ($p < 0.01$) in the young group compared to the elderly group. (b) ABIDE-I dataset: violin plots corresponding to 425 typically developing (TD) individuals and 395 individuals with autism spectrum disorder (ASD) across three global measures. Average persistent entropy is significantly higher ($p < 0.001$) in the ASD group than the TD group; however, average $L^1$-norm and $L^2$-norm are significantly lower ($p < 0.001$) in the ASD group.

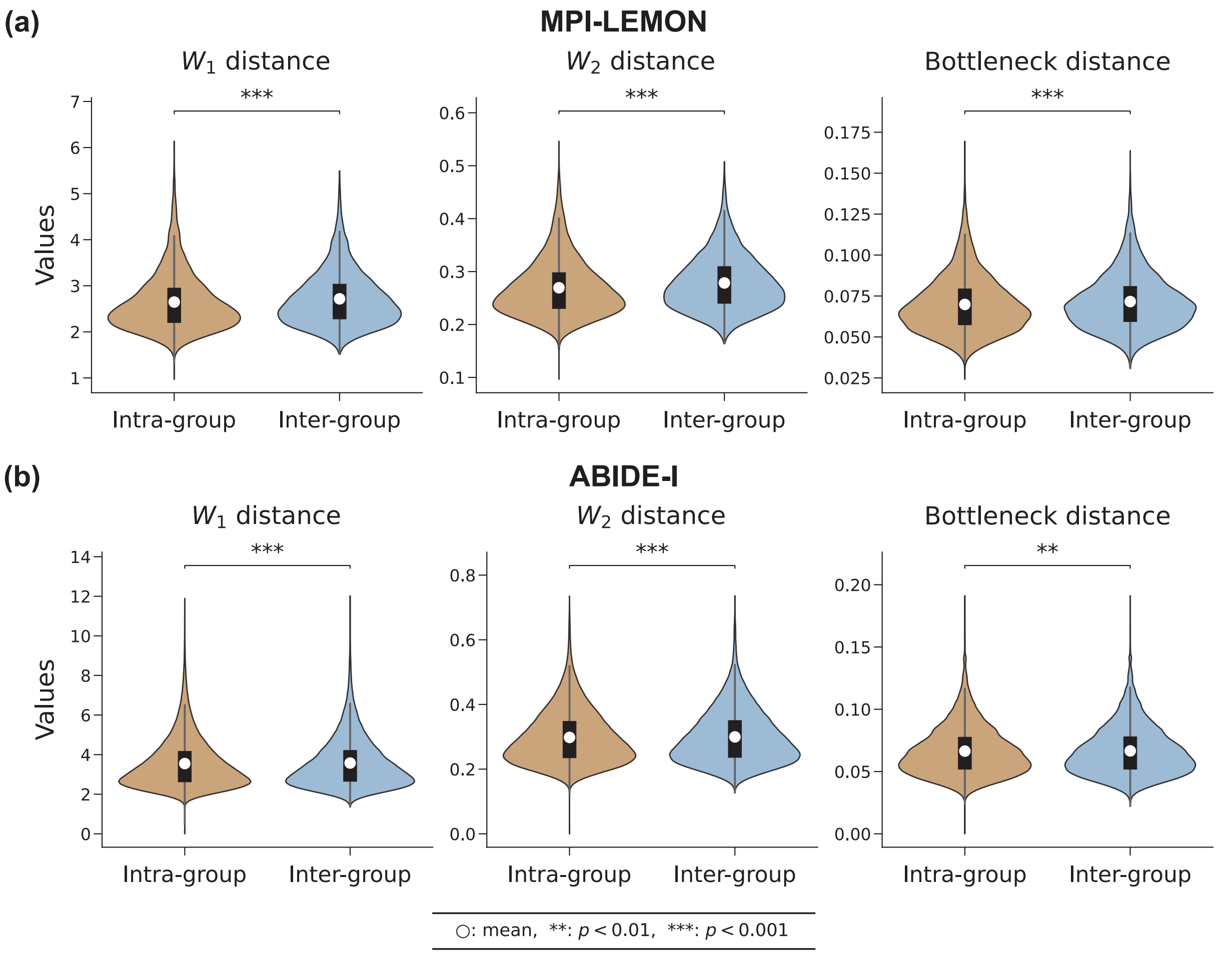


FIG. S2. **Brain-wide differences between the intra-group and inter-group distances of persistent diagrams as identified by three global measures: 1-Wasserstein ($W_1$), 2-Wasserstein ($W_2$), and bottleneck distances, considering only the positive correlations of the functional connectivity matrices.** (a) MPI-LEMON dataset: violin plots corresponding to intra-group (young-young or elderly-elderly pairs) and inter-group (young-elderly pairs) distances across three measures. The mean values of inter-group distances are significantly higher ($p < 0.001$) than the intra-group distances. (b) ABIDE-I dataset: violin plots corresponding to intra-group (ASD-ASD or TD-TD pairs) and inter-group (ASD-TD pairs) distances across three global measures. The mean values of inter-group distances are significantly higher ($p < 0.01$) than the intra-group distances.

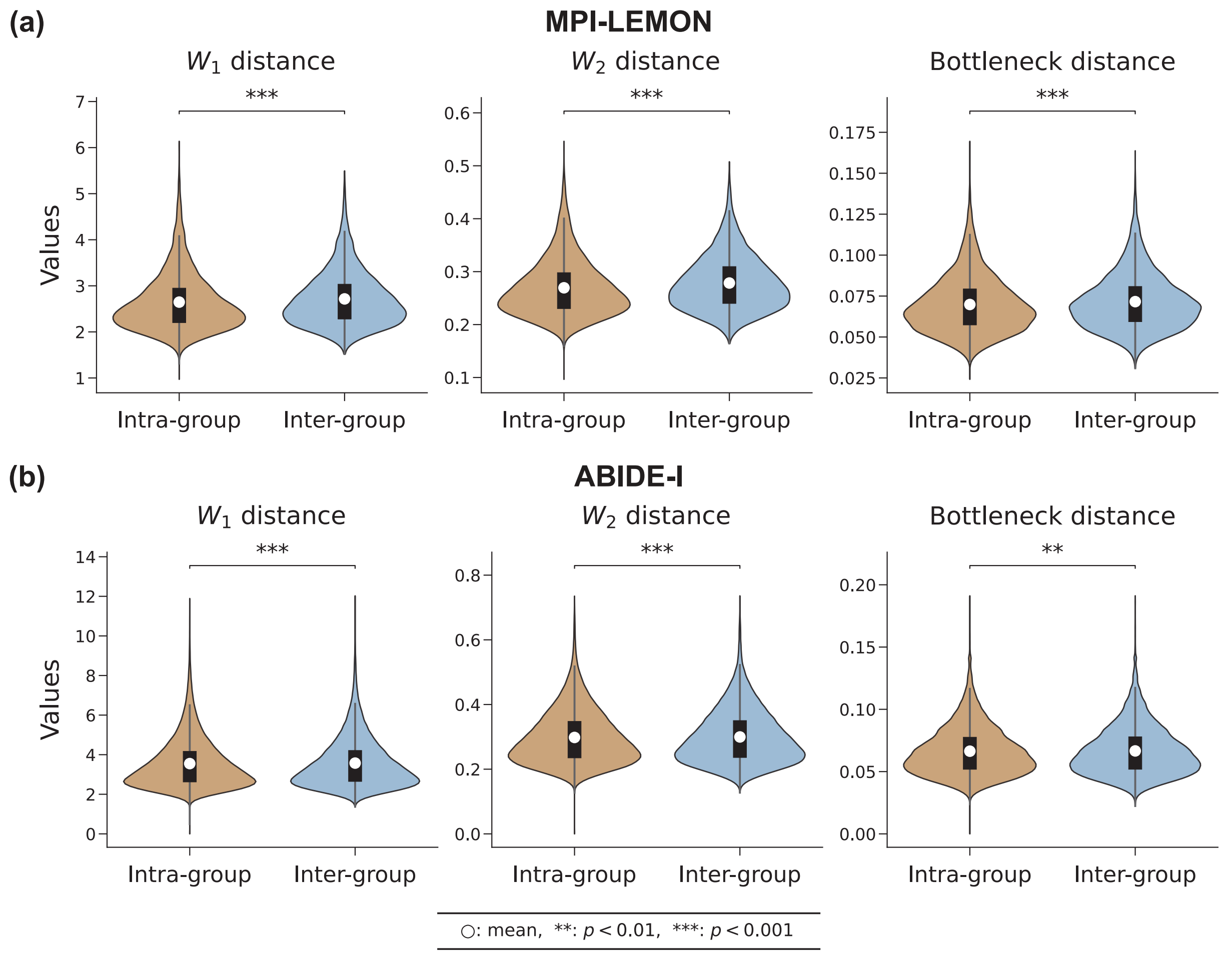


FIG. S3. **Brain-wide differences between the intra-group and inter-group distances of persistent diagrams as identified by three global measures: 1-Wasserstein ($W_1$), 2-Wasserstein ($W_2$), and bottleneck distances, considering all the correlations of the functional connectivity matrices.** (a) MPI-LEMON dataset: violin plots corresponding to intra-group (young-young or elderly-elderly pairs) and inter-group (young-elderly pairs) distances across three measures. The mean values of inter-group distances are significantly higher ($p < 0.001$) than the intra-group distances. (b) ABIDE-I dataset: violin plots corresponding to intra-group (ASD-ASD or TD-TD pairs) and inter-group (ASD-TD pairs) distances across three global measures. The mean values of inter-group distances are significantly higher ($p < 0.01$) than the intra-group distances.

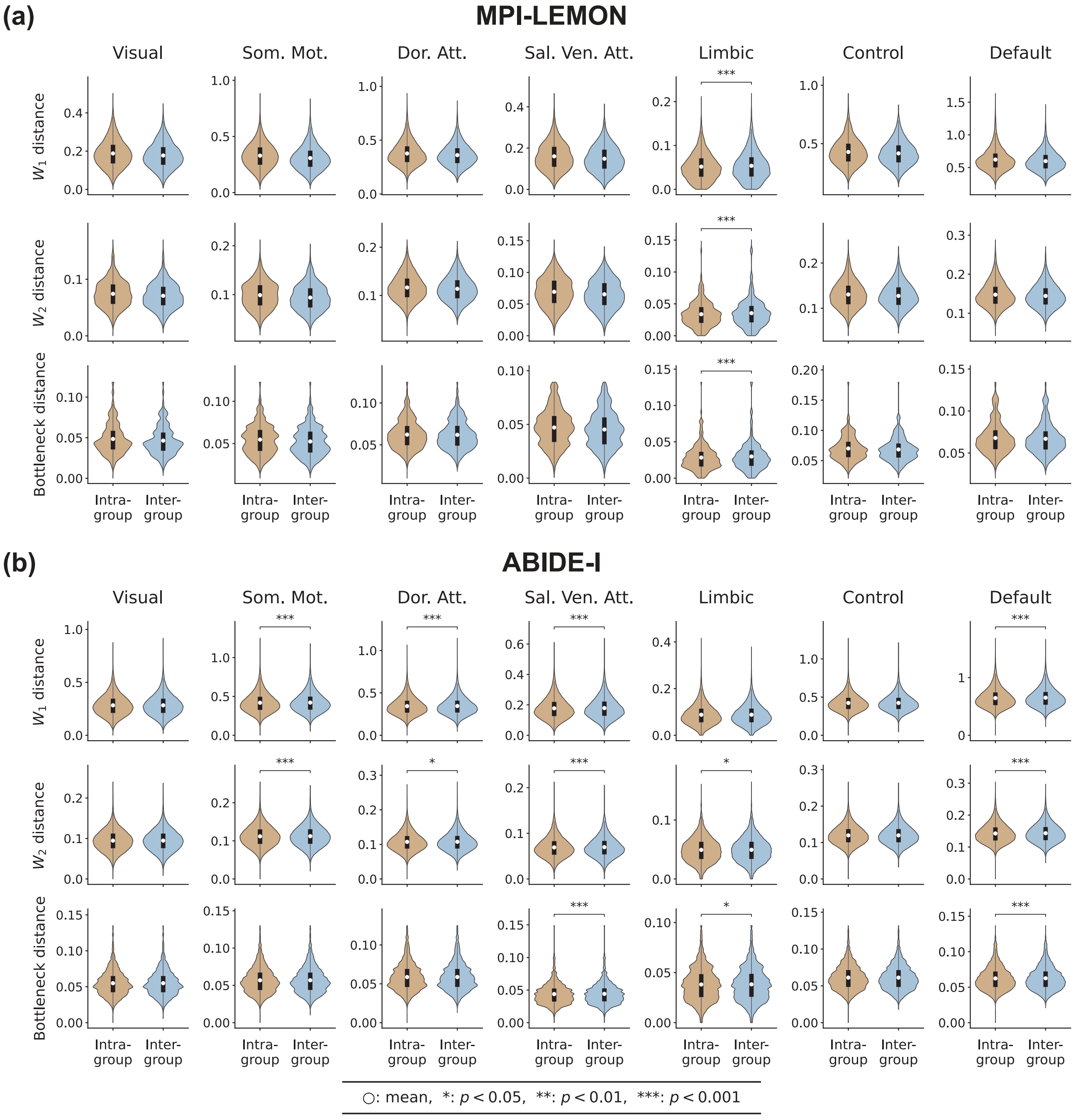


FIG. S4. **RSN-level differences between the intra-group and inter-group distances of the persistent diagrams as identified by three global measures: 1-Wasserstein ($W_1$), 2-Wasserstein ($W_2$), and bottleneck distances, considering only the positive correlations of the functional connectivity matrices.** Each row corresponds to a given measure, and each column corresponds to a given RSN. (a) MPI-LEMON dataset: violin plots corresponding to intra-group (young-young or elderly-elderly pairs) and inter-group (young-elderly pairs) distances. Inter-group distances are significantly higher ($p < 0.001$) than the intra-group distances only for the limbic network corresponding to all three measures. (b) ABIDE-I dataset: violin plots corresponding to intra-group (ASD-ASD or TD-TD pairs) and inter-group (ASD-TD pairs) distances. For the somatomotor (Som. Mot.) and dorsal attention (Dor. Att.) networks, the inter-group distances are significantly higher ($p < 0.05$) than the intra-group distances for 1-Wasserstein and 2-Wasserstein distances. In the salience/ventral attention (Sal. Ven. Att.) and default networks, inter-group distances are significantly higher ($p < 0.001$) than the intra-group distances for the three measures: 1-Wasserstein, 2-Wasserstein and bottleneck distances. In the limbic network, inter-group distances are significantly higher ($p < 0.05$) than the intra-group distances for 2-Wasserstein and bottleneck distances.

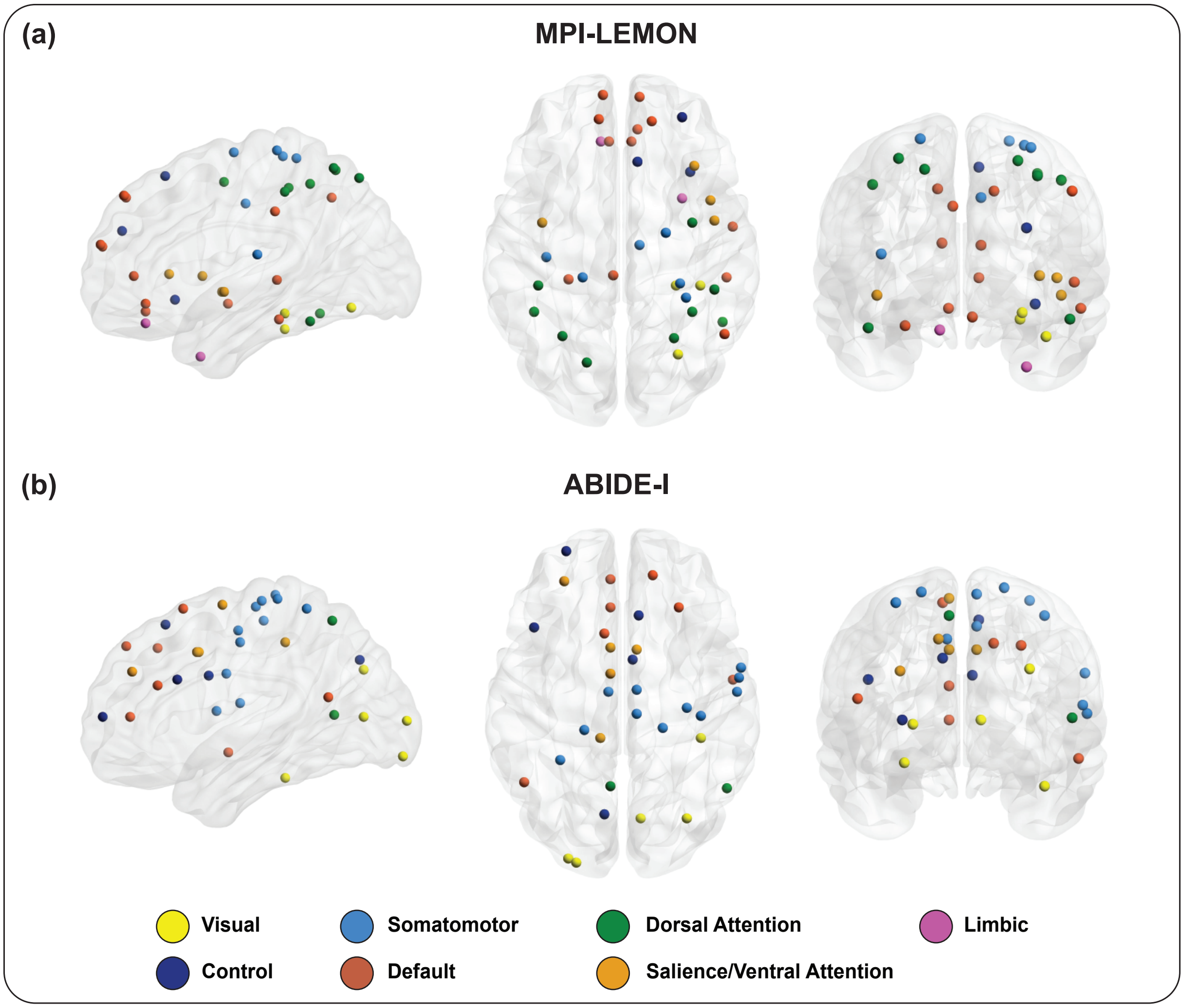


FIG. S5. **Visual representation of brain regions with significant between-group differences in node frequency ($p < 0.05$, FDR-corrected)** (a) MPI-LEMON dataset: 39 regions with significant differences in node frequency between the healthy young and healthy elderly groups. For every region, young participants show higher node frequency compared to the elderly participants, except one region RH_Limbic_TempPole_1. (b) ABIDE-I dataset: 35 regions with significant differences in node frequency between the autism spectrum disorder (ASD) and typically developing (TD) groups. All regions reveal increased node frequency for ASD participants relative to TD participants. Each brain region is assigned to one of the seven resting-state networks (RSNs) as defined by the Schaefer atlas. The regions are colored according to their respective RSNs, as detailed in the figure legend. The visualization was generated using BrainNet Viewer. Supplementary Table S3 lists the significantly different ROIs identified via node frequency across both datasets.

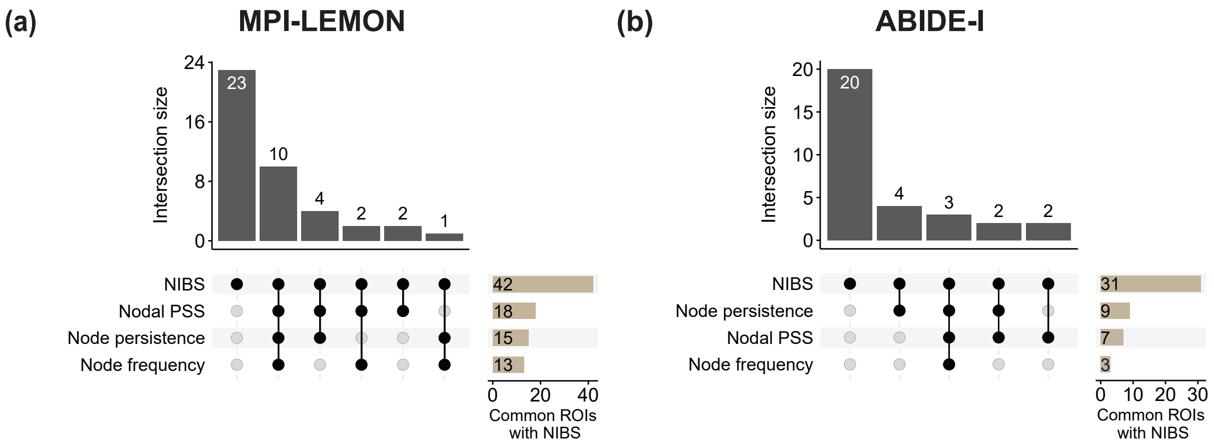


FIG. S6. **UpSet plots illustrating the intersection of ROIs identified via non-invasive brain stimulation (NIBS), and local persistent homology-based measures from RSN-to-local level analysis.** Each subplot highlights the combinations of node-based measures, namely node persistence, node frequency, and nodal PSS, with a focus on NIBS-identified regions. The connected cells in the matrix indicate the sets of ROIs involved in each intersection, and the bars aligned with the columns represent the size of each intersection, providing a visual representation of how the sets overlap, particularly emphasizing the role of NIBS in these intersections. The right horizontal bars indicate the cardinality of the intersection of the set of ROIs identified via NIBS and corresponding local persistent homology-based measures. (a) MPI-LEMON dataset: within NIBS, 19 of 42 ROIs are identified considering all three measures collectively, and 10 ROIs are identified when all three are considered simultaneously. Two ROIs are uniquely identified using nodal PSS. (b) ABIDE-I dataset: within the 31 ROIs in NIBS, 11 ROIs are identified considering all three measures collectively, and three ROIs are common across all the measures. Four and two ROIs are uniquely identified considering node persistence and nodal PSS, respectively.
